# MAVS signaling is required for clearance of chikungunya heart infection and prevention of chronic inflammation in vascular tissue

**DOI:** 10.1101/2022.05.27.493768

**Authors:** Maria G. Noval, Sophie N. Spector, Eric Bartnicki, Franco Izzo, Navneet Narula, Stephen T. Yeung, Payal Damani-Yokota, M. Zahidunnabi Dewan, Valeria Mezzano, Bruno A. Rodriguez-Rodriguez, Cynthia Loomis, Kamal M. Khanna, Kenneth A. Stapleford

**Affiliations:** Department of Microbiology, New York University Grossman School of Medicine, New York, NY, USA; New York Genome Center, New York, NY, USA; Division of Hematology and Medical Oncology, Department of Medicine and Meyer Cancer Center, Weill Cornell Medicine, New York, NY, USA; Department of Pathology, New York University Grossman School of Medicine, New York, NY, USA; Division of Advanced Research Technologies, New York University Grossman School of Medicine, New York, NY, USA; Perlmutter Cancer Center, New York University Grossman School of Medicine, New York, NY, USA

**Author notes:** Corresponding authors: Kenneth A. Stapleford. and Maria G. Noval. Lead contact: Kenneth A. Stapleford.

**Keywords:** Chikungunya virus, pathogenesis, heart, myocarditis, vasculitis, type-I interferon, MAVS

## Abstract

Chikungunya virus (CHIKV) infection has been associated with severe cardiac manifestations, yet, how CHIKV infection leads to heart disease remains unknown. Here, we leveraged both mouse models and human primary cells to define the mechanisms of CHIKV heart infection. We found that CHIKV actively replicates in cardiac fibroblasts and is cleared without significant tissue damage through the induction of a local type-I interferon response from both infected and non-infected cardiac cells. Importantly, signaling through the mitochondrial antiviral-signaling protein (MAVS) is required for viral clearance from the heart. In the absence of MAVS, persistent infection leads to focal myocarditis and major vessel vasculitis persisting for up to 60 days post-infection, suggesting CHIKV can lead to vascular inflammation and potential long-lasting cardiovascular complications. This study provides a model of CHIKV cardiac infection and mechanistic insight into CHIKV-induced heart disease, underscoring the importance of monitoring cardiac function in patients with CHIKV infections.

## INTRODUCTION

Arthropod-borne viruses (arboviruses) such as Zika virus, dengue virus, and chikungunya virus (CHIKV) are associated with the development of cardiomyopathies in patients^1-6^. CHIKV-associated cardiac manifestations have been reported in more than 15 countries worldwide^5,7^. Cardiac complications occur within the first week after symptoms onset^5^, and include symptoms ranging from arrhythmias, atrial fibrillation, abnormal echocardiograms and electrocardiograms^8^, reduction of the ejection fraction^6^, myocarditis^8-10^, to heart failure, and death^9,11-13^. Autopsies of individuals that succumbed to CHIKV infection reveal the presence of CHIKV antigen in the cardiac tissue^12,14^ and immune cell infiltration indicative of viral myocarditis, as well as signs of cardiac edema, endocarditis, necrosis, and cardiac congestion^9^. Indeed, more than 20% of the CHIKV-related mortality cases have been associated with cardiac complications^9,12,15^. Despite most of cardiac complications being observed in older adults and individuals with comorbidities^11,16^, cardiac manifestations associated with CHIKV infections have been reported in young individuals, including children and infants without known comorbidities^6,8,9^.

While previous studies reported a link between CHIKV and infection of cardiac tissue in animal models^17-21^, several questions remain unanswered: (i) is cardiac tissue a direct target of CHIKV infection and a site of active replication in immunocompetent hosts?; (ii) what are the mechanisms of CHIKV-cardiac tissue interaction?; (iii) how does the heart respond to CHIKV infection?; and (iv) how does CHIKV infection lead to heart disease?.

Here, using an immunocompetent mouse model of CHIKV infection as well as human primary cardiac cells, we demonstrate that CHIKV directly infects and actively replicates within cardiac tissue, with cardiac fibroblasts being the main cellular target of CHIKV infection. We show that CHIKV infects the myocardium, valves, atrium, and vessels, and that viral RNA persists longer in tissue containing atrium and vessels in comparison to the myocardium. We show that CHIKV heart infection is rapidly cleared without signs of tissue inflammation or cardiac tissue damage in immunocompetent mice coinciding with a local type I interferon (IFN-I) response. Indeed, cardiac tissue infection induces a local IFN-I response in both infected and non-infected cardiac cells, and the loss of IFN-I signaling results in increased CHIKV infectious particles, virus spreading, and apoptosis in cardiac tissue. We demonstrate that the IFN-I response is essential to control CHIKV infection in human primary cardiac fibroblasts. In addition, we found that signaling through the mitochondrial antiviral-signaling protein (MAVS), a central hub for signal transduction initiated by cytosolic pattern recognition receptors (PRRs) RIG-I or MDA5, is required for efficient CHIKV clearance, with viral particles being detected up to 10 days post-infection (dpi) in *Mavs*^*-/-*^ mice. Importantly, we found that the persistence of CHIKV infection in cardiac tissue of *Mavs*^*-/-*^ mice leads to cardiac tissue damage characterized by CD3^+^ and CD11b^+^ infiltrates in myocardium and atrium as well as in the major vessels attached to the base of the heart, such as the aorta (Ao) and the pulmonary artery (PA). Interestingly, viral myocarditis was detected at 10 and 15 dpi, while major vessel vasculitis involving the Ao and PA persisted up to 60 dpi, suggesting that infection of cardiac tissue by CHIKV can lead to vascular inflammation and potential long-lasting cardiovascular complications.

Overall, our results show that a robust IFN-I response and MAVS signaling are required to control CHIKV infection in the heart and prevent focal myocarditis as well as Ao and PA vasculitis. Altogether, this study provides mechanistic insight into how CHIKV can lead to cardiac damage, underscoring the importance of monitoring cardiac function in patients with CHIKV infections and paving the way towards the identification of risk factors associated to the development of CHIKV-induced cardiac complications.

## RESULTS

### CHIKV actively replicates in cardiac tissue and targets cardiac fibroblasts

While the presence of CHIKV viral products have been detected in cardiac tissue of infected animals^17-21^, whether CHIKV is infecting and replicating within cardiac cells remained elusive. To define if cardiac tissue is a direct target of CHIKV infection in immunocompetent mice, we infected mice either subcutaneously through footpad injection, intravenously via tail vein injection, or via natural transmission by mosquito bite. To account for the potential confounder of viremia (viral particles coming from the bloodstream rather than the tissue), we perfused hearts to clear the tissue from residual blood and associated virus. We quantified CHIKV infectious virus by median tissue culture infectious dose (TCID_50_) and CHIKV RNA by RT-qPCR. We were able to detect CHIKV genomes and infectious particles from perfused hearts in infected mice regardless of the inoculation route **(Fig. 1a and Extended Fig. 1a)**, supporting cardiac tissue as a *bona fide* target of CHIKV infection. To evaluate if the CHIKV viral particles detected in the heart were actively replicating, we measured the ratio between CHIKV nsp4 and E1 transcripts by RT-qPCR as a surrogate of genomic and sub-genomic RNA, respectively, at 2, 5 and 10 dpi. As a non-replicative control, we infected mice with heat-inactivated (HI) CHIKV. While no active replication was detected for the HI control, CHIKV active replication in the heart was detected for both subcutaneous and intravenous inoculation routes **(Fig. 1b)**. For intravenously infected mice active replication was detected at 2 dpi (R [Ratio E1/nsp4] = 3.80, p = 0.04) and 5 dpi (R = 8.11, p = 0.008), while for subcutaneously infected mice active replication was detected at 5 dpi (R = 3.88, p = 0.008). The difference in kinetics between inoculation routes is consistent with the requirement of subcutaneous inoculation to reach viremia in order to disseminate and infect cardiac tissue, while the intravenous route bypasses this stage of the infection. These results demonstrate that cardiac tissue is a direct target of CHIKV infection in immunocompetent mice.

**Fig. 1.**
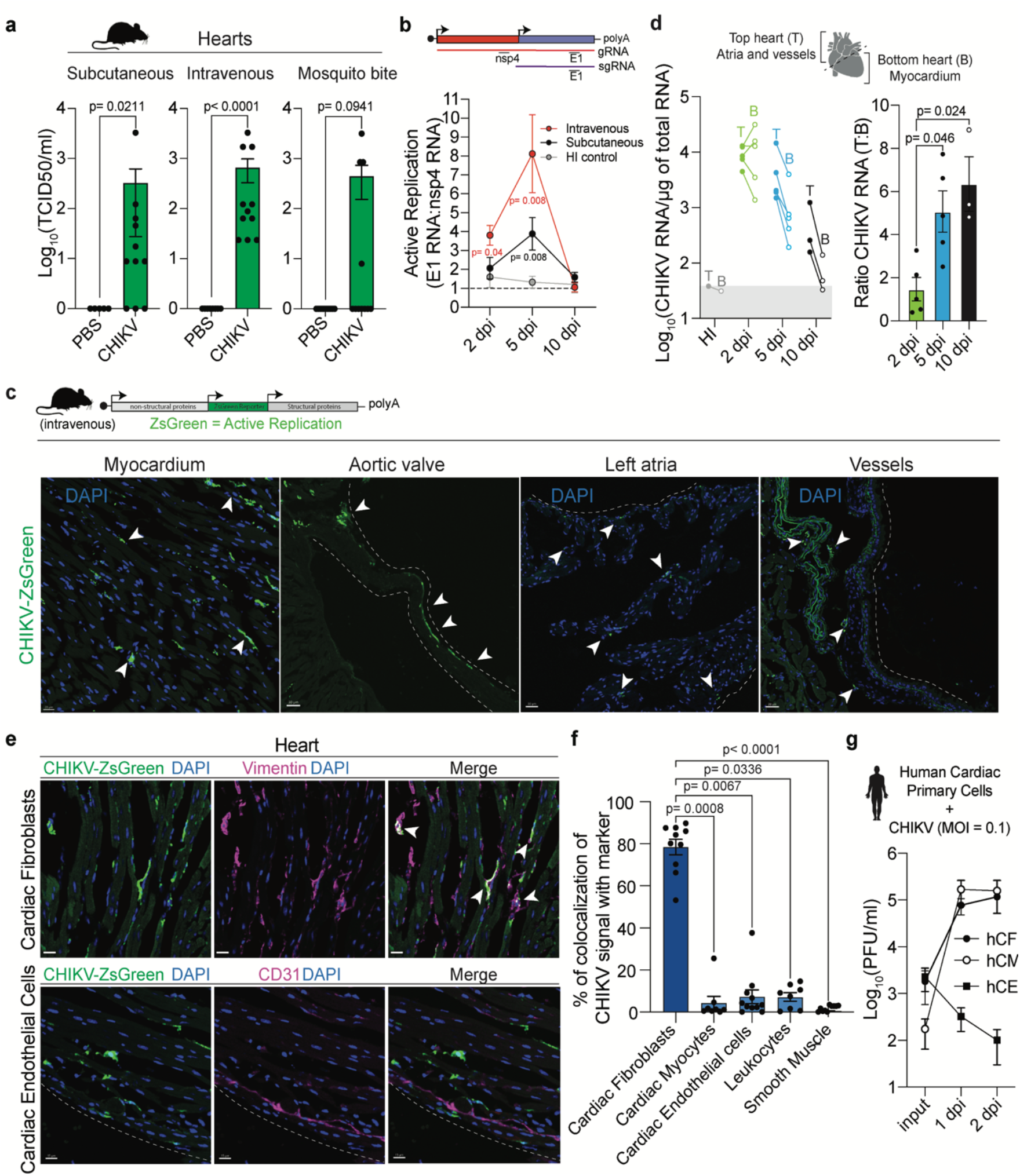
CHIKV actively replicates in cardiac tissue and targets cardiac fibroblasts. **a**. Six-week-old C57BL/6 mice were inoculated with 1E5 PFU of CHIKV or mock-infected subcutaneously, intravenously or via natural transmission through a mosquito bite. Mice were harvested at 2 dpi for subcutaneous and intravenous infection routes, and at 2 and 3 dpi for mice infected through mosquito bite. Infectious particles were quantified by TCID_50_. N=5-11 mice/group. **b**. Mice were infected either by subcutaneous (black dots) or intravenous routes (red dots), and tissue was harvested at 2, 5, and 10 dpi. The ratio of E1 viral RNA to nsp4 viral RNA was determined. Dotted line indicates the E1 RNA:nsp4 RNA ratio (R) of 1. R > 1 indicates active viral replication. N=4-8/group. **c**. Representative images of infected myocardium, aortic valve, left atria, and vessels. Green channel: CHIKV-infected cells. Blue channel: DAPI. White arrows: CHIKV-infected cells. Scale= 20-30 μm. **d**. Mice were inoculated with 1E5 PFU of CHIKV or HI virus intravenously. Hearts were divided into the top and the bottom sections and CHIKV RNA levels were quantified by RT-qPCR. Left panel: CHIKV RNA levels for top and bottom hearts. Right panel: ratio of viral RNA levels between top and bottom heart sections. N=3-5 mice/group. The gray box represents the background RNA levels quantified for HI control at 5 dpi. **e**. Representative fluorescence microscopy images of ventricular sections of CHIKV-infected hearts stained with cardiac fibroblast markers (top panel) or cardiac endothelial cell markers (bottom panel). Scale= 15 μm. **f**. Co-localization analysis between CHIKV-infected cells and different cell type markers. Data are represented as the percentage of the total green signal (infected cells) overlapping with the indicated markers. Colocalization analysis was done in three to four fields for N=3 mice. **g**. CHIKV multi-cycle growth curves in human primary cardiac fibroblast (hCF, back dot), human primary cardiac myocytes (hCM, open dots) and human primary cardiac endothelial cells (hCE, square). Data represent the average of two independent repetitions in technical duplicates. Data represents two to

To determine the localization of CHIKV infection within cardiac tissue, we used a reporter CHIKV that expresses the ZsGreen fluorescent protein (CHIKV-ZsGreen) exclusively under active viral replication^17^. Mice were infected with CHIKV-ZsGreen and cardiac tissue was harvested at 2 dpi. We observed multiple ZsGreen positive foci corresponding with CHIKV-infected cells for mice infected both intravenously **(Fig 1c)** as well as subcutaneously **(Extended Data Fig 1b)**. CHIKV can infect different structures of the cardiac tissue, with presence of infected cells within the myocardium, aortic valves, atrium, and vessels **(Fig 1c)**. With the objective of measuring the differences in the level of infection across cardiac tissue, we physically separated the heart into: (i) top section (containing atria and enriched in vessels attached to the base of the heart, such as ascending thoracic aorta) and (ii) bottom section (enrich in the myocardium); and evaluated the kinetics of CHIKV infection.

To address direct infection of cardiac tissue and to bypass the variability associated with differences in dissemination, we used the intravenous infection route unless stated otherwise. We found that while CHIKV can replicate efficiently in both top and bottom sections of the heart at 2 dpi (R =1.47; **Fig.1d left panel**), CHIKV RNA genomes persist for longer times in the top section containing the vessels and atria (5 dpi: R=5.07, and 10 dpi: R=6.35, **Fig 1d, left panel)**. Indeed, we found that the top section of the heart contains higher amounts of viral genomes relative to the bottom section (**Fig 1d, right panel**). These results support that different structures of the cardiac tissue are not equally susceptible to CHIKV infection, and that the section containing large vessels sustain higher CHIKV viral load.

To identify the specific cell types infected by CHIKV in cardiac tissue, hearts were harvested at 2 dpi, fixed, and processed to obtain serial cryosections followed by staining with markers for specific cardiac cell types: CD31 (endothelial cells), CD45 (leukocytes), αSMA (smooth muscle cells, pericytes, and myofibroblasts), cTnT (cardiomyocytes), and vimentin. Of note, vimentin is expressed at high levels in cardiac fibroblasts^22^ but can also be expressed in smooth muscle cells, endothelial cells, and macrophages^23-25^. Therefore, we identified cardiac fibroblasts cells as vimentin^+^/αSMA^-^/CD31^-^/CD45^-^populations, a combination that we orthogonally validated via single-cell RNA expression from the *Tabula Muris*^26^ **(Extended Data Fig. 1c**). By using fluorescence microscopy and co-localization analysis with the different cell type markers, we found that the 78.5% of CHIKV-ZsGreen signal colocalized with vimentin^+^/αSMA^-^ cells **(Fig. 1e-f, and Extended Data Fig. 1d and 1e)**. This result, together with the reduced colocalization between CHIKV-ZsGreen signal and markers for endothelial cells (CD31^+^; 7.3%), leukocytes (CD45^+^; 7.2%), smooth muscle/pericytes/myofibroblasts (αSMA^+^; 1%), and cardiomyocytes (cTnT^+^; 4.4%; **Fig. 1f; Extended Data Fig. 1e)** demonstrate that the cardiac fibroblasts are the primary target of CHIKV infection of cardiac tissue *in vivo*.

Finally, we sought to determine whether human cardiac cells show similar susceptibilities to CHIKV-infection than those observed in mice. For that purpose, we evaluated the capacity of CHIKV to infect human primary cardiac cells *in vitro*. We infected human primary cardiac fibroblasts (hCF), human cardiac myocytes (hCM) and human cardiac microvasculature endothelial cells (hCE) at an MOI of 0.1 and measured the production of infectious particles. We found that CHIKV can efficiently infect hCF (**Fig. 1g, black circles**), but no hCE **(Fig. 1g, squares)**, supporting our *in vivo* observations in mice. Interestingly, hCM in culture produced CHIKV infectious particles at similar levels than the observed for hCFs **(Fig. 1g, open circles)**. However, careful interpretation of this result must be taken, since hCMs in culture loose specific features that can potentially impact susceptibility to CHIKV infection. For example, while cardiac myocytes in tissue are characterized for being multinucleated and quiescent *in vivo*, the *in vitro* culture revert these phenotypes, resulting in mono-nucleated and proliferating cells^27^.

Altogether, our results demonstrate that CHIKV actively replicates in cardiac tissue with cardiac fibroblasts being the primary cellular target of CHIKV infection in immunocompetent mice.

### Cardiac tissue clears CHIKV infection without inducing cardiac damage in immunocompetent mice

To evaluate the consequences of CHIKV replication in cardiac tissue, we measured the kinetics of CHIKV infection in the heart, as well as cardiac damage, apoptosis and inflammation in cardiac tissue. Significant levels of CHIKV RNA were detected in the heart of intravenously infected mice as early as 12 hours post-infection (hpi) with a peak at 2 dpi, and the infection was cleared out to background detection levels by 9 dpi **(Fig. 2a, left panel)**. Infectious particles follow a similar trend, with a peak of infectious particle production between 1 and 2 dpi, and no detectable levels by 9 dpi **(Fig. 2a, right panel)**. Interestingly, albeit with higher CHIKV RNA levels, the viral replication kinetics follow a similar temporal pattern when measured in calf muscle, a primary organ of CHIKV replication **(Extended Data Fig. 2a)**. Therefore, these results demonstrate that CHIKV infection is efficiently cleared of cardiac tissue in immunocompetent mice.

**Fig. 2.**
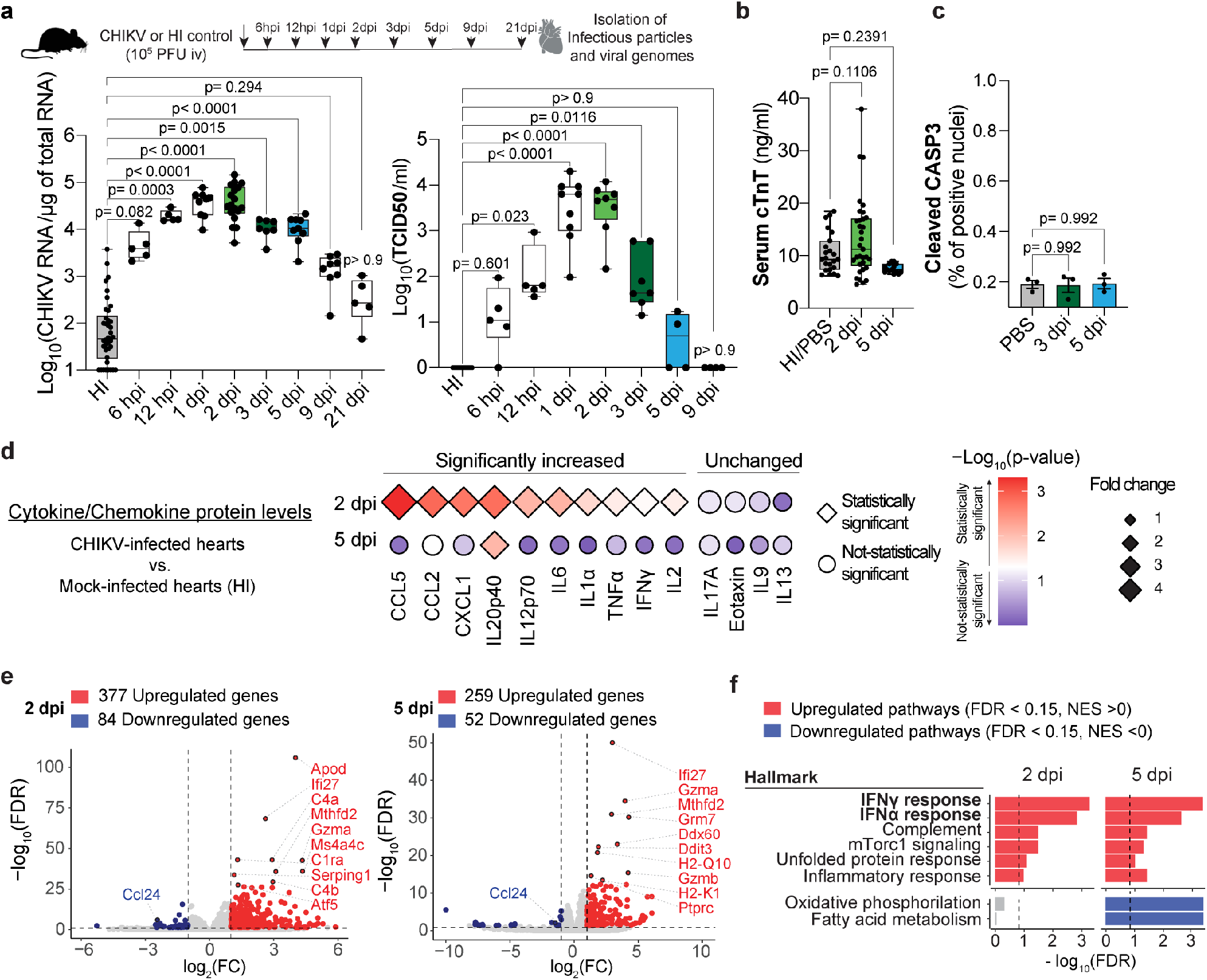
Cardiac tissue clears CHIKV infection without inducing cardiac damage in immunocompetent mice. **a**. CHIKV heart infection kinetics. Six-week-old C57BL/6 mice were inoculated intravenously with 1E5 PFU of CHIKV or mock-infected (HI). Mice were harvested at the indicated time points. Animals were perfused with 30 ml of cold PBS, and hearts were isolated for downstream processing. Upper panel: Schematic representation of the experimental design. Bottom panel: CHIKV viral genomes or CHIKV infectious particles determined by RT-qPCR or TCID_50_ in BHK-21 cells, respectively. N=5-9 mice/group. **b**. Serum levels of cardiac troponin-T measured by ELISA. Mock: HI or PBS. N=14-30 mice/group **c**. Determination of cardiac tissue apoptosis by IHC. Tissue was stained for cleaved CASP3 (see **Extended Data Fig.2b** for positive control). Quantifications of cleaved CASP3 nuclei versus total nuclei were performed in whole ventricular sections (LV, RV, and septum) as described in materials and methods using the software Visiopharm^R^. Each dot represents one individual mouse. N=3 mice/group. **d**. Plot showing the relative protein levels of proinflammatory cytokines and chemokines between CHIKV-versus HI control-infected cardiac tissue homogenates at 2 and 5 dpi. Relative expression is represented in log2 fold change diamond size. Statistical significance is expressed as a color scale (-Log_10_ adjusted p-value). Statistically significant values are represented as diamonds, while no significant changes are represented as circles. Samples were measured using the Bio-Plex Pro Mouse Cytokine 23-plex Assay. N=5-6 mice/group. **e**. Volcano plots from differential expression analysis at 2 dpi (left panel) and 5 dpi (right panel). Red and blue indicate upregulated (FDR < 0.15 and log_2_(FC) > 1) and downregulated genes (FDR < 0.15 and log_2_(FC) < -1), respectively. Top-10 differentially expressed genes are indicated. N=4 mice/group. **f**. GSEA pathway enrichment analysis for Hallmark datasets showing top upregulated (FDR < 0.15 and normalized enrichment score (NES) > 0, red bars) and downregulated (FDR < 0.15 and NES < 0, blue bars) pathways at 2 dpi and 5 dpi. Mock: HI control. N=4 mice/group.

Elevated cardiac troponins have been observed in individuals with CHIKV-associated cardiac manifestations^6,7,13,16^. To define whether CHIKV can lead to cardiac damage in mice, we evaluated the serum levels of cardiac troponin-T (cTnT) at 2 and 5 dpi by ELISA. We observed no significant differences in levels of cardiac troponin-T in serum between infected and PBS or HI controls (**Fig. 2b)**, suggesting that CHIKV replication does not induce damage to cardiac myocytes. Indeed, no sign of cardiac tissue apoptosis was observed at 3 or 5 dpi **(Fig. 2c**, positive staining control in **Extended Data Figure 2b)**. Thus, no cardiac damage is observed in WT mice upon CHIKV infection and clearance.

Despite no cardiac damage was observed upon CHIKV in WT mice, we found that CHIKV infection stimulates the protein production of proinflammatory cytokine and chemokine in cardiac tissue (CCL2, CCL5, CCL3, TNFα, IL6, IFNγ, IL2, KC, IL1α, IL12p70, and IL12p40) at 2 dpi. However, IL12p40 is the only cytokine that remains upregulated at 5 dpi **(Fig. 2d; Extended Data Fig. 2c-d)**, indicating that CHIKV infection is inducing a transient inflammatory state in cardiac tissue. Histopathological analysis of hematoxylin-eosin (H&E) staining of four-chamber view sections of infected and mock-infected at 3, 5, 10, and 15 dpi **(Extended Data Table 1)** revealed no visible mononuclear cell infiltrates in ventricles, septum, or atria of CHIKV-infected hearts. Altogether, these results demonstrate that cardiac tissue can efficiently clear CHIKV infection without significant cardiac damage in immunocompetent mice.

### Local IFN-I response in cardiac tissue is essential to control CHIKV infection and prevent tissue damage

To understand how the heart rapidly clears CHIKV infection without cardiac tissue damage in WT mice, we measured the transcriptional response of CHIKV-infected heart homogenates by bulk RNA-seq. We used CHIKV-infected mice at 2 dpi (where we detect both CHIKV RNA and CHIKV infectious particles) or at 5 dpi (where we detect little to no CHIKV infectious particles, but CHIKV RNA levels are detected; **Fig. 2a**). To identify changes derived only from active CHIKV replication, we used the HI virus as a control. Principal component analysis (PCA) showed that biological replicates grouped by treatment **(Extended Data Fig. 3a)**. A total of 377 upregulated genes and 84 downregulated genes were detected at 2 dpi, and 259 upregulated genes as well as 52 downregulated genes at 5 dpi **(Fig. 2e, Extended Data Fig. 3b-c; Extended Data Table 2-3)**. The top-10 upregulated genes on both days include genes associated with protection of cardiac tissue damage and mitochondria metabolism (*Apod, Atf5, Mthfd2*)^28,29^, IFN-I response (*Ifi27, Ddx60, Serping1*), cytokine signaling (*H2-K1*), and complement cascade (*C4a, C4b, C1ra*) **(Fig. 2e)**. In line with these observations, GSEA pathway enrichment analysis for Reactome and Hallmark datasets show upregulation (FDR < 0.15 and normalized enrichment score [NES] > 0) of several pathways associated with innate immunity, adaptive immunity, IFN signaling **(Fig. 2f, Extended Data Fig. 3c, and Table 4)**. Downregulated pathways (FDR < 0.15 and NES < 0) in infected heart homogenates at 5 dpi include metabolic pathways such as metabolism of amino acids, respiratory electron transport, and oxidative phosphorylation, among others **(Extended Data Fig 3c**).

The induction of IFN-I signaling pathways in non-hematopoietic cells is essential to control CHIKV infection in mammals^30,31^. We found that the IFN-I pathway is significantly induced in CHIKV-infected hearts at both 2 and 5 dpi (**Fig. 3a, and Extended Data Fig. 3e**). Particularly, we observed upregulation of interferon stimulated genes (ISGs; e.g. *Isg15, Ifitm3, Mx1, Mx2, Ifit1, Ifit2, Ifit3, ifi27, Ddx58*, and *Ddx60*, among others) as well as genes involved in the IFN-I signaling pathway (e.g. *Irf7, Stat1, Stat2*; **Extended Data Table 2-3**). While the levels of serum IFNα and IFNβ were elevated at 24 hpi, no changes in local levels of IFNα and IFNβ were observed in infected hearts at 12 or 24 hpi (**Extended Data Fig. 3f**). This suggests that circulating IFN-I is a contributor to the local IFN-I response elicited by cardiac tissue upon CHIKV infection.

**Fig. 3.**
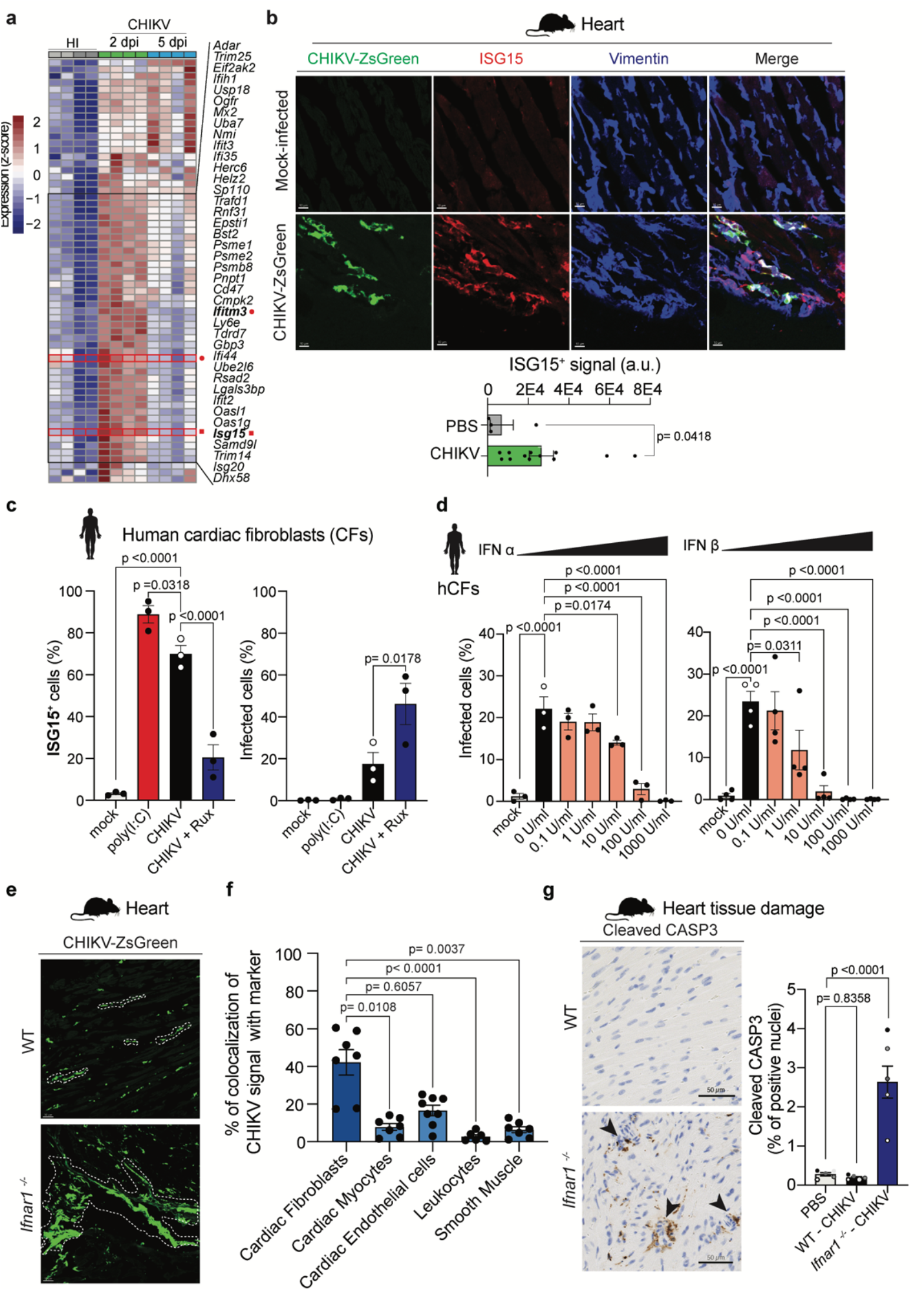
Local IFN-I response in cardiac tissue control CHIKV infection and prevent tissue damage. **a**. Heat map showing the differentially expressed genes (FDR < 0.15) from the Hallmark IFNα response pathway for CHIKV-infected and HI-infected hearts. **b**. Top panel: representative fluorescence microscopy images of ventricular sections of CHIKV-infected hearts and PBS control stained with anti-ISG15 and anti-vimentin antibodies. Scale = 10 μm. Bottom panel: Quantification of ISG15^+^ signal/field. Analysis was done on independent sections for N=3 mice/group. **c**. Dependence on IFN-I signaling in hCFs during CHIKV infection. hCFs were pretreated as indicated in the figure and infected with CHIKV-ZsGreen at an MOI of 0.25. IFN-I response (% of ISG15^+^ cells) and Infection (% of infected cells) were determined by high-content microscopy. **d**. IFN-I pretreatment experiments in hCFs. hCFs were pre-treated with different concentrations of recombinant IFN-α (right panel) and IFN-β (left panel) for 24 h and then infected with CHIKV-ZsGreen at an MOI of 0.25 for 24 h. Infection (% of infected cells was determined by high-content microscopy). **(c-d)**. Data represent three to four independent repetitions and each dot represents the average of two technical replicates. **e**. Representative fluorescence microscopy images of CHIKV-ZsGreen WT or *Ifnar1*^-/-^ infected hearts at 1 dpi. White shapes highlight CHIKV infection spreading. Scale= 30 μm. **f**. Percentage of the total of infected cells (green signal) overlapping with the indicated makers (**See Extended Data Fig 1c-d**). Co-localization analysis was done in three to four independent sections for N=2 mice. **g**. Left panel: Apoptosis staining was measured by IHC against cleaved CASP3. Scale= 50 μm. Arrows highlight apoptotic foci. Right panel: Quantifications of cleaved CASP3 nuclei versus total nuclei were performed in whole ventricular sections (LV, RV, and septum) using Visiopharm^R^. Each dot represents one individual mouse. N=5 mice/group. P values were calculated using Mann-Whitney **(b)** or Kruskal-Wallis test (**c-d, f-g)**.

To gain insight into whether the IFN-I response observed by RNA-seq was locally mediated by infected cardiac fibroblasts, we measured the local protein production of IFITM3 and ISG15, both ISGs known to restrict CHIKV infection^32,33^, and found that both were upregulated at 2 dpi in CHIKV-infected hearts **(Fig. 3a, and Extended Data Fig. 3d**). While IFITM3 is expressed at basal levels in the absence of interferon induction, its expression is highly induced by either IFN-I or interferon-γ^34^. We found that that 95% of CHIKV-infected cells are positive for the IFITM3 marker. However, in the absence of the IFN-I receptor (interferon a/b receptor-deficient mice (*Ifnar1*^*-/-*^) only 24.6% of CHIKV-infected cells colocalize with the IFITM3 signal (**Extended Data Fig. 4a**), demonstrating that the observed response is partially dependent on IFN-I. Local production of ISG15 and IFITM3 proteins were detected in cardiac tissue upon CHIKV infection in both infected and non-infected vimentin^+^ cells (**Fig. 3b. and Extended Data Fig. 4a**), suggesting that cardiac fibroblast among other vimentin^+^ cells are the major IFN-I responders in infected cardiac tissue.

Next, we sought to determine the importance of IFN-I signaling in the infection of human cardiac fibroblasts (hCFs) by CHIKV. We evaluated this by: (i) measuring the capacity of hCFs to generate a robust IFN-I response after poly(I:C) stimulation and CHIKV infection; (ii) measuring the susceptibility of these cells to CHIKV infection in the absence of IFN-I signaling using the JAK1/JAK2 inhibitor ruxolitinib (Rux); and (iii) inducing a refractory state upon increasing doses of IFN-I pre-treatments. Indeed, we found that hCFs efficiently respond to poly(I:C) stimulation, resulting in ∼ 80% of the monolayer expressing ISG15 (**Fig. 3c, left panel**). In agreement with our *in vivo* data, we found that infected hCFs responded to CHIKV infection by producing significant levels of ISG15 **(Fig. 3c, left panel)**. In addition, Rux treatment significantly increased the percentage of CHIKV infected cells (**Fig. 3c, right panel**), and correlated with a reduction of ISG15 production **(Fig. 3c, left panel**). Furthermore, we found that exogenous addition of either IFNα or IFNβ prevented CHIKV infection in a dose response manner, with IFNβ showing increased blocking capacity (**Fig 3d**). These results support that IFN-I signaling is critical in controlling CHIKV infection in human cardiac fibroblasts.

Finally, we evaluated the direct contribution of the IFN-I response in controlling CHIKV heart infection and preventing tissue damage by using *Ifnar1*^*-/-*^ mice. As expected, we observed that in the absence of IFN-I signaling, CHIKV spreading and infectious particle production were increased in cardiac tissue of *Ifnar1*^*-/-*^ mice compared to immunocompetent WT mice **(Fig. 3e, and Extended Data Fig. 4b)**. This increase in infectious particle production was also observed in the calf muscle, a primary site of CHIKV infection, but not observed in a non-CHIKV-target organ such as the pancreas **(Extended Data Fig. 4b)**. While cardiac fibroblasts (Vimentin^+^/aSMA^-^; 42%) still represent the primary target of CHIKV infection in cardiac tissue in *Ifnar1*^*-/-*^ mice, we observed an increased proportion of endothelial infected cells (CD31^+^; 16.5%, **Fig. 3f, and Extended Data Fig. 3c)** relative to WT (**Fig. 1f**). Intriguingly, in *Ifnar1*^*-/-*^ mice only 7.8% of the infected cells colocalized with cTnT marker (**Fig. 3f, and Extended Data Fig. 3c)**, suggesting that even in the absence of IFN-I response, cardiac myocytes do not represent a target of CHIKV infection in mice. To determine whether the observed increased infection in *Ifnar1*^*-/-*^ mice results in cardiac damage even in the absence of cardiac myocyte infection, we assessed cardiac tissue apoptosis by cleaved CASP3 staining at 1 dpi. We observed that while infection of WT mice showed similar levels of cleaved CASP3 signal in comparison to mock-infected control, *Ifnar1*^*-/-*^ mice showed a significantly increased level of cleaved CASP3 within the myocardium (**Fig. 3g)**, demonstrating that unrestricted CHIKV infection can lead to cardiac tissue damage. Altogether, these results demonstrate that a local IFN-I response is essential to control CHIKV infection in the heart and prevent cardiac tissue damage.

### MAVS signaling is required for viral clearance in CHIKV-infected hearts

IFN-I signaling is triggered upon recognition of CHIKV RNA via either cytosolic (RIG-I/MDA5) or endosomal receptors (TLR3/7)^35^. To gain insight into the mechanisms of how cardiac tissue clears CHIKV infection, we took advantage of mice deficient in major innate immune signaling components including MYD88, TRIF, or MAVS. In addition, to address for the contribution of the NLRP3 inflammasome in CHIKV clearance^36,37^ we used CASPASE 1/11 deficient mice. We infected each mouse intravenously with CHIKV and evaluated viral burden in cardiac tissue at 2, 5, and 7 dpi. Interestingly, while we found no significant differences in the infection kinetics between *Myd88* ^*-/-*^, *Trif* ^*Lps2/Lps2*^ or *Casp1/11*^*-/-*^ mice compared to WT mice, infectious particles were detectable up to 7 dpi for *Mavs*^*-/-*^ mice **(Fig. 4a)**. Indeed, we found that the heterozygote deletion of MAVS results in a phenotype similar to WT, demonstrating that the complete loss of MAVS is required for persistence CHIKV infection **(Extended Data Fig. 5a)**. These results suggested that MAVS signaling is contributing to the clearance of CHIKV infection from infected hearts.

**Fig. 4.**
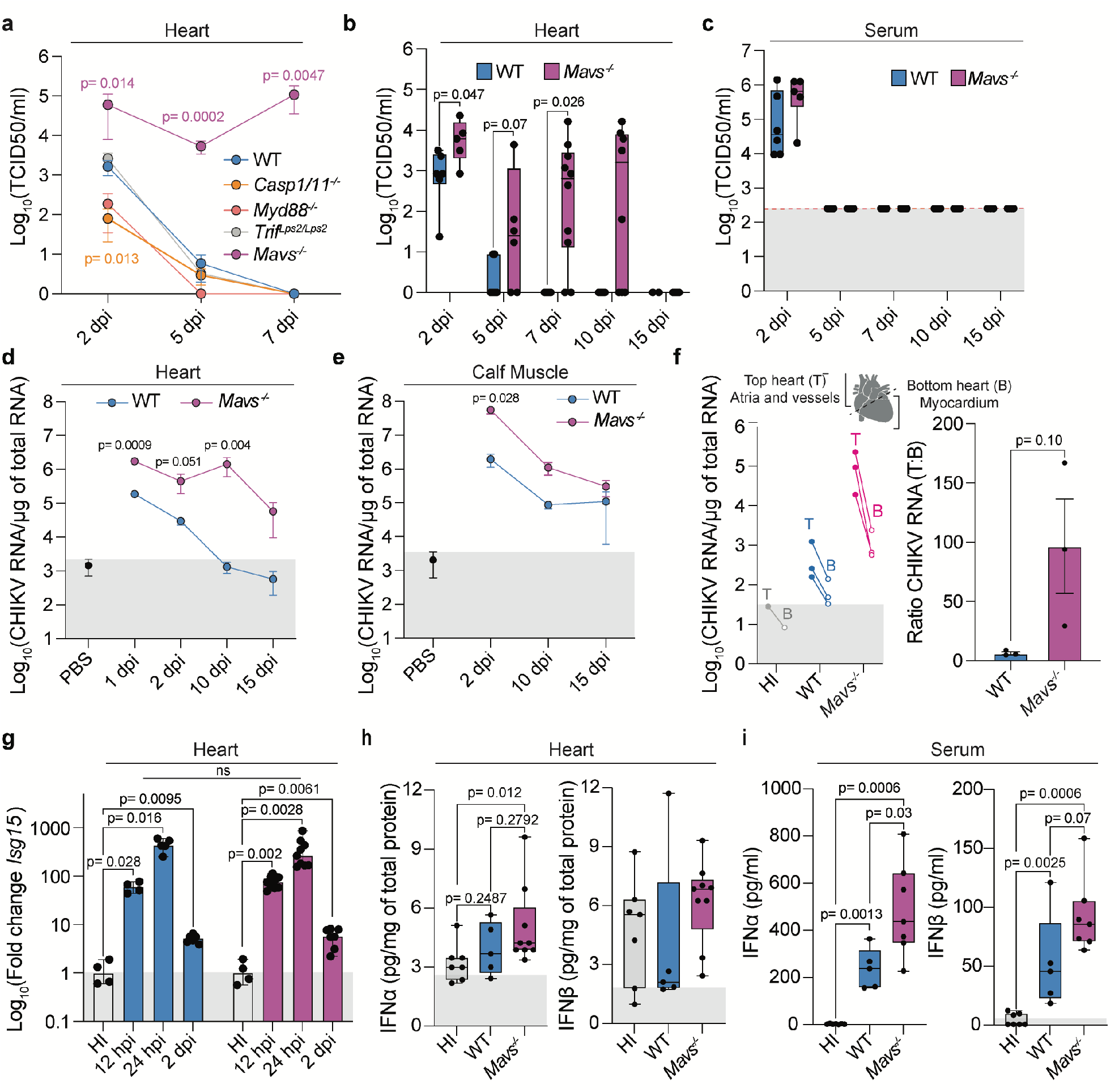
MAVS signaling is required for CHIKV clearance in infected hearts. **a**. Differential susceptibilities to CHIKV heart infection between *MyD88*^-/-^, *Trif*^*Lps2/Lps2*^, *Casp1/11*^-/-^, *Mavs*^-/-^, and WT mice at 2, 5, and 7 dpi. Mice were infected intravenously with 1E5 PFU of CHIKV. CHIKV infectious particles were determined by TCID_50_ in BHK-21 cells. N=3-10 mice/group. **b-c** CHIKV heart infection kinetics in *Mavs*^-/-^ and WT mice. Mice were infected intravenously with 1E5 PFU of CHIKV. CHIKV infectious particles from heart homogenates **(b)** and from serum **(c)** were determined by TCID_50_ in BHK-21. N=4-10 mice/group, with the exception of WT at 15 dpi (N=2). Gray box represents the limit of detection in serum. **d-e**. CHIKV RNA genomes from the heart **(d)** and from calf muscle homogenates **(e)** were determined by RT-qPCR. N= 3-10 mice/group. The gray box represents the background RNA levels determined for PBS controls. **f**. *Mavs*^-/-^ and WT mice were infected intravenously with 1E5 PFU of CHIKV and harvested at 10 dpi. Left panel: hearts were divided into top and bottom sections and CHIKV RNA levels were quantified by RT-qPCR. Data set for WT at 10 dpi is the same from **Fig 1c**. N=3 mice/group. Right panel: ratio of viral RNA levels between top and bottom heart sections. N= 3-5 mice/group. **g**. Dynamics of *Isg15* expression levels in *Mavs*^-/-^ and WT infected hearts. N=4-9 mice/group. Gray boxes indicate a fold change of 1. **h-i**. Protein levels of IFN-α and INF-β in cardiac tissue homogenates **(h)** and serum **(i)** at 24 hpi. N=5-7 mice/group. The gray box represents values in which the concentration of the analyte was determined by extrapolation. P values were calculated using Mann-Whitney (**a-i)**.

Next, we expanded our analysis up to 15 dpi and we found that infectious particles can be detected up to 10 dpi **(Fig. 4b)**, while infectious particles in serum were cleared by 5 dpi for both *Mavs*^*-/-*^ and WT mice **(Fig. 4c)**. In line with this, CHIKV-ZsGreen infected cells were detected up to 10 dpi in both myocardial region as well as in vessels in *Mavs*^*-/-*^ mice **(Extended Data Fig. 5b-c)**. To evaluate whether this deficiency in viral clearance from cardiac tissue is observed in other tissues we evaluated the levels of CHIKV RNA in calf muscle and heart at 2, 10, and 15 dpi for both WT and *Mavs*^*-/-*^ mice. We found that the kinetics of CHIKV clearance from the calf muscle is different from cardiac tissue. For cardiac tissue, we observed a progressive clearance in the RNA levels of WT mice, for *Mavs*^*-/-*^ mice we observed that the levels of viral RNA are maintained up to 10 dpi, and evidence of RNA decay is observed by 15 dpi **(Fig. 4d)**. However, in calf muscle, both *Mavs*^*-/-*^ and WT mice showed a progressive reduction of viral RNA over time, and at 10 dpi no significant differences in CHIKV RNA were observed **(Fig. 4e)**. Interestingly, when we evaluated the CHIKV RNA levels in the top section or bottom section of the heart at 10 dpi, we observed higher viral RNA levels in top sections relative to the bottom sections **(Fig. 4f)**. These results are in agreement with our observation in WT animals **(Fig. 1d)**, and supports that heart sections enriched in atria and/or vessels are more prone to sustained infection over time. Collectively, these results demonstrated that sensing through MAVS is required for clearance of CHIKV infection from cardiac tissue.

Of note, we found that *Mavs*^*-/-*^ infected hearts induced similar levels of *Isg15* mRNA compared to WT (**Fig. 4g)**. Prompted by this result, we measured the local (cardiac tissue homogenate) and systemic (serum) production of IFNα or IFNβ at 24 hpi, which corresponds to the peak of induction of *Isg15* in cardiac tissue **(Fig. 4g)**. We found that while the local IFN-I production is low in cardiac tissue **(Fig. 4h)**, both WT and *Mavs*^*-/-*^ mice produce significant systemic levels of both IFNα and IFNβ **(Fig. 4i)**. Surprisingly, the systemic levels of both IFNα and IFNβ were higher in *Mavs*^*-/-*^ mice compared to WT **(Fig. 4i)**, which correlates with the significantly higher viral loads observed in *Mavs*^*-/-*^ **(Fig. 4b and 4d-e)**. These data suggest that the delay in CHIKV clearance cannot be entirely explained by a reduced IFN-I induction associated with *Mavs*^*-/-*^.

### *Mavs*^*-/-*^ mice develop myocarditis and major vessel vasculitis upon CHIKV infection

Next, we sought to determine whether the persistence of CHIKV-heart infection in *Mavs*^*-/-*^ mice can lead to cardiac tissue damage. We infected *Mavs*^*-/-*^, *Mavs*^*+/-*^, and WT mice intravenously with CHIKV and harvested the heart for histopathological analysis at 10 dpi **(Fig.5a, upper panel)**. As expected, we didn’t observe any signs of inflammation in the myocardial tissue for neither *Mavs*^*+/-*^ nor WT mice **(Fig.5a-b and Extended Data Table 1)**. However, 53% of CHIKV-infected *Mavs*^*-/-*^ mice (7 out of 13 mice) developed focal myocarditis at 10 dpi **(Fig.5a-b and Extended Data Table 1)**. Strikingly, we found that all of the CHIKV-infected *Mavs*^*-/-*^ mice (13 out of 13) have transmural inflammation of the vessel wall in the major vessels attached to the base of the heart (e.g., aorta, pulmonary artery) corresponding to major vessel vasculitis **(Fig. 5a and c, and Extended Data Table 1)**. Of note, a fraction of WT and the *Mavs*^*+/-*^ mice showed minimal signs of inflammation in the vessels corresponding to endothelialitis, characterized by mild inflammation of the intima layer of the vessel **(Fig. 5a, bottom panel)**. Altogether, these results demonstrate that the persistence of CHIKV infection in cardiac tissue can lead to inflammatory infiltrates leading to focal myocarditis and major vessel cell vasculitis.

**Fig. 5.**
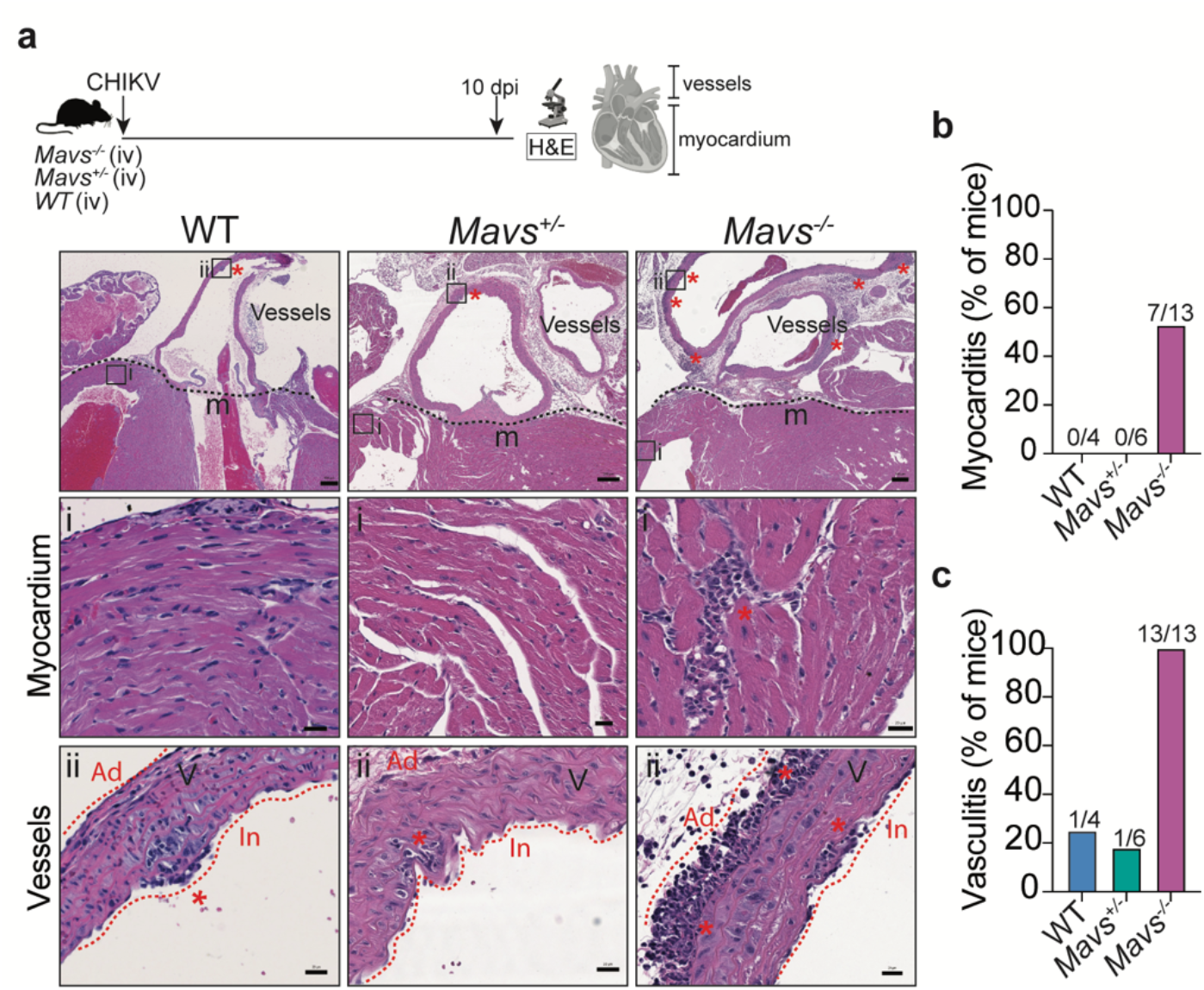
*Mavs*^*-/-*^ mice develop myocarditis and major vessel vasculitis upon CHIKV infection. **a**. Upper panel: Schematic representation of the experimental design. Bottom panel: Representative H&E for CHIKV-infected WT, *Mavs*^*+/-*^ and *Mavs*^*-/-*^ hearts at 10 dpi. Scale = 200 μm. The black square indicates the section selected. Lower Panel: Representative selections of the myocardium and major vessels attached to the base of the heart. Asterisks highlight the presence of immune cell infiltrates. Adventitia (Ad) and intima (In) layers are indicated in red. Scale = 20 μm. **b**. Quantification of the number of mice with positive signs of myocarditis. Data represent the percentage of mice with any sign of myocarditis over the total amount of mice analyzed. N=4-13 mice/group. **c**. Quantification of the number of mice with major vessel vasculitis. Data represent the percentage of mice with any sign of vasculitis over the total amount of mice analyzed. N=4-13 mice/group. (Related to **Extended Data Table 1**).

### Chronic inflammation in myocardium and vasculature in CHIKV-infected *Mavs*^*-/-*^ mice persist for several months

Given the focal myocarditis and the major vessel vasculitis observed at 10 dpi in *Mavs*^*-/-*^ mice, we asked whether this inflammation can result in chronic inflammation potentially leading to long-term tissue damage. For that purpose, *Mavs*^*-/-*^ mice were infected intravenously or subcutaneously with CHIKV, and cardiac tissue was collected for histopathological analysis at 2 weeks, 1 month, or 2 months post-infection. Tissue was stained with H&E, and mononuclear cell infiltrates were evaluated for CD3 and CD11b markers by IHC. We found that a fraction of *Mavs*^*-/-*^ mice developed focal and multifocal myocarditis accentuated around the vessels in mice infected by either intravenous or subcutaneous inoculation routes (**Fig. 6a-b and Extended Data Table 1**). These inflammatory foci were characterized by CD11b^+^, and CD3^+^ cell infiltrates and persisted for up to 60 dpi or up to 31 dpi in mice infected intravenously or subcutaneously, respectively **(Fig.6b, and Extended Data Fig. 6a-b and 6e)**. Interestingly, we did not detect changes in the serum levels of cardiac Troponin T **(Fig. 6c)**, suggesting that despite myocarditis there is no significant cardiac myocyte damage in this model. Strikingly, major vasculitis characterized by transmural CD3^+^ andCD11b^+^ infiltrates was detected up to 60 dpi in infected *Mavs*^*-/-*^ mice independently of the inoculation route (**Fig. 6d; Extended Fig. 6c-d, and Extended Data Table 1)**.

**Fig. 6.**
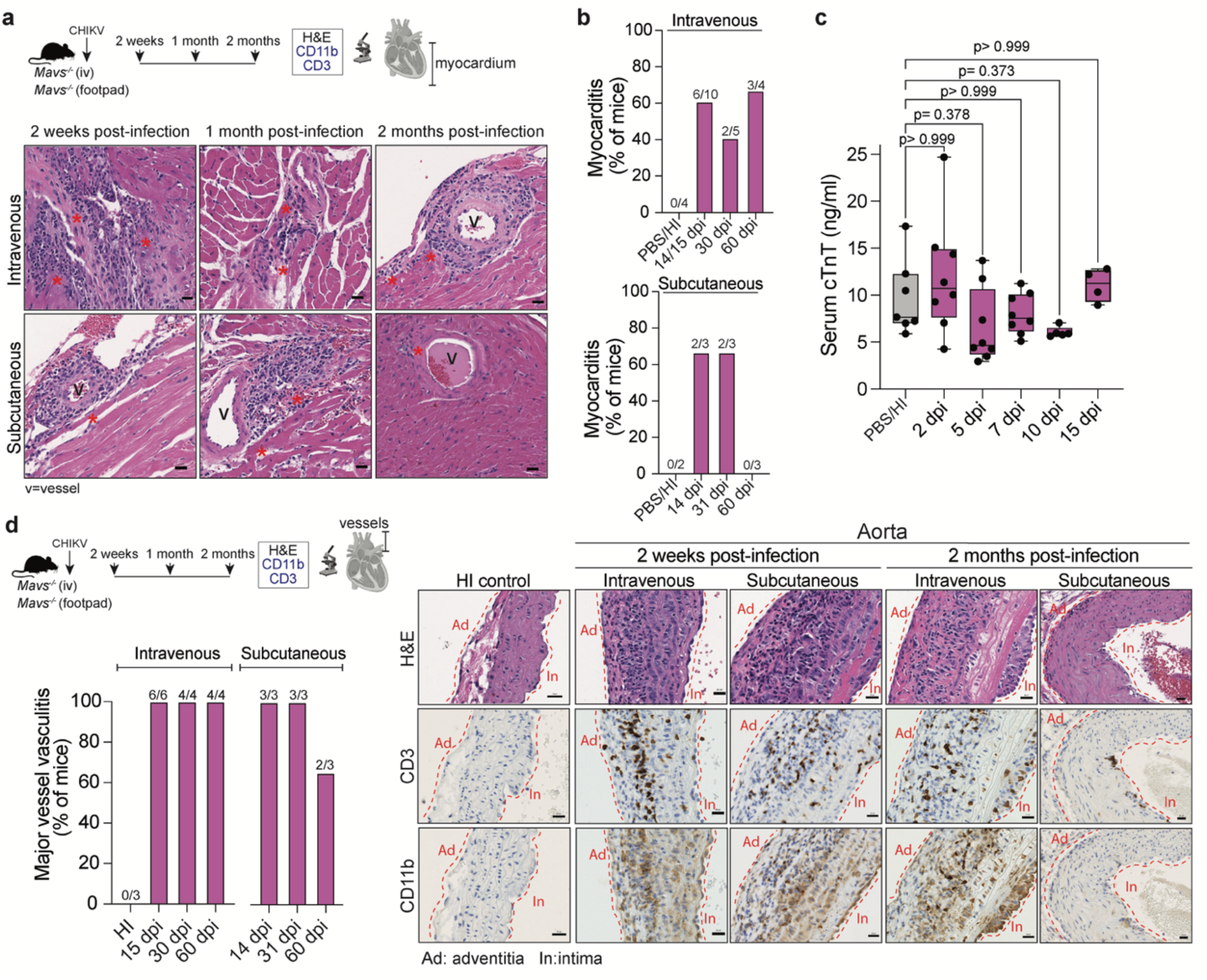
CHIKV infected *Mavs*^*-/-*^ mice develop chronic inflammation in myocardium and vasculature. **a**. Upper panel: Schematic representation of the experimental design. Bottom panel: Representative H&E sections from the myocardium of subcutaneous or intravenously inoculated CHIKV-infected *Mavs*^*-/-*^ mice at 2 weeks, 1 month, or 2 months post-infection. Asterisks indicate the presence of immune cell infiltrates. Vessels within the myocardial region are indicated. Scale = 20 μm. **b**. Quantification of the number of mice with positive signs of myocarditis for both intravenous and subcutaneous inoculation routes. Data represent the percentage of mice with any sign of myocarditis over the total amount of mice analyzed. N=2-10 mice/group. **c**. Serum levels of cardiac troponin-T measured by ELISA. N=4-8 mice/group. **d** Schematic representation of the experimental design. Left panel: quantification of the number of mice with major vessel vasculitis. Data represent the percentage of mice with any sign of vasculitis over the total amount of mice analyzed. N=3-6 mice/group. Right panel: representative H&E and CD3/CD11b IHC sections from the vessels attached to the base of the heart of subcutaneous or intravenously inoculated CHIKV-infected *Mavs*^*-/-*^ mice at 2 weeks and 2 months post-infection. Scale= 20 μm. Related to **Extended Data Table 1 and Extended Fig 6**. P values were calculated using multiple Mann-Whitney tests **(c)**.

Of note, cardiac tissue of an infected *Mavs*^*-/-*^ mice that succumbed to CHIKV infection at 15 dpi (**Extended Data Table 5**), showed substantial tissue damage featured by patchy myocyte dystrophic calcification affecting single myocytes, and focal myocarditis composed by inflammatory infiltrates of CD11b^+^ and CD3^+^ cells, with no signs of fibrosis **(Extended Data Fig. 7)**. Altogether, these results demonstrate that persistence of CHIKV infection in cardiac tissue leads to chronic inflammation characterized by focal or multifocal myocarditis and major vessel vasculitis.

## Discussion

In this study, we defined mechanisms implicated in CHIKV cardiac tissue infection and identified host factors involved in the development of cardiac tissue damage. Here, we demonstrated: (i) that cardiac fibroblasts are a direct target of CHIKV infection in an immunocompetent host; (ii) that cardiac tissue elicits a local IFN-I response required for CHIKV clearance and preventing tissue damage; (iii) that in the absence of MAVS signaling CHIKV persists in infected hearts, and (iv) that failure to clear CHIKV in *Mavs*^*-/-*^ mice leads to focal myocarditis and chronic major vessel vasculitis detectable up to 60 dpi.

Previous studies, including our own, reported the presence of infectious particles or viral RNA in non-perfused cardiac tissue of CHIKV-infected mice^19-21^. However, due to the lack of perfusion and high levels of CHIKV in circulation, whether cardiac tissue is a direct target of CHIKV infection remained elusive. Here, by including a perfusion step, and by using orthogonal experimental approaches, we demonstrated that CHIKV actively replicates in the cardiac tissue of immunocompetent mice regardless of the infection route. Moreover, we showed that CHIKV infects cardiac fibroblasts and that its infection follows a discrete and focalized pattern within ventricles, atria, valves, and the perivascular region. We found that primary human cardiac fibroblasts are susceptible and permissive to CHIKV. Indeed, CHIKV antigen in cardiac fibroblasts was previously identified in human necropsies of individuals who died from CHIKV^12^. Therefore, our results in murine models are consistent with the tropism observed in human necropsies.

We demonstrated that in WT mice, CHIKV is cleared from cardiac tissue within the first week of infection without causing tissue damage. Our RNA-seq data revealed that infected cardiac tissue shows upregulation of innate immunity, adaptive immunity, and IFN-I signaling pathways, similar to what has been described for footpad and lymph node CHIKV-infected tissue^38^. In addition, we found that genes previously associated with protection of the cardiac tissue and antioxidant metabolism (*Apod, Aft5, Fgf21, and Mthfd2*) were upregulated at 2 dpi^39,40^. *Apod* is a cardioprotective gene whose expression is induced in mouse hearts during myocardial infarction and has been proposed to protect the heart against ischemic pathway at 5 dpi **(Extended Data Fig. 3c and g-i; Extended Data Table 4)**. However, we did not observe signs of cardiac tissue damage by histopathology (**Extended Data Table S1**), suggesting that CHIKV infection is triggering transcriptional programs involved in cardiac dysfunction that are not damage^28^. In addition, *Apod* has been reported to be the most significantly upregulated gene in SARS-CoV-2-infected cardiac tissue in fatal COVID-19 cases^41^ and has been found upregulated in a cluster of cardiac fibroblasts during Coxsackievirus B3 infection^42^. Further studies are required to determine the specific role of *Apod* in cardioprotection during CHIKV infection.

Intriguingly, using the KEGG datasets we observed upregulation of the viral myocarditis pathway and downregulation of the cardiac tissue contractility sufficient to result in overt phenotypes of cardiac damage.

We found that CHIKV-infected hearts display a significant yet transient production of proinflammatory chemokines and cytokines, and induction of IFITM3 and ISG15 proteins in both CHIKV-infected and not-infected vimentin^+^ cardiac cells. Interestingly, it has been shown that upon TLRs or RIG-I-like receptor stimulation, human cardiac fibroblasts can produce proinflammatory cytokines and induce an antiviral immune response^43^. Our study confirms the critical role of IFN-I signaling in controlling CHIKV infection in human cardiac fibroblasts. In the context of an immunocompetent host, the infected cardiac tissue mounts a robust and local IFN-I response resolving CHIKV infection within days, with little to no cardiac damage. This observation provides an explanation of why CHIKV cardiac manifestation is not a pervasive outcome in CHIKV-infected individuals.

In line with this, a report using 12-days-old KO mice for a protein implicated in the production of IFN-β during CHIKV infection (GADD34) revealed a significant increase in viral particles in the heart and development of myocarditis compared to WT^21^. Similar results were described for IFITM3 KO mice during influenza virus infection^44^. Therefore, the risk of cardiac complications in CHIKV-infected patients could be associated with genetic and non-genetic factors (such as co-infections, diet, among others) leading to a reduced IFN-I response. Single nucleotide polymorphisms (SNPs) in TLR3 found in patients with enteroviral myocarditis are associated with a decreased TLR3 mediated signaling and shown an impaired IFN-I response after Coxsackievirus B3 infection^45^. Thus, SNPs in IFN-I and *MAVS* signaling genes could underlie an increased susceptibility to cardiac complications in CHIKV-infected patients. SNPs in the coding region of the human *MAVS* can result in impaired function^46,47^. In addition, recent studies suggested a link between age and decreased sensing through *MAVS*^48^, which could be a contributing factor to the observed correlation between age and CHIKV severity^5,15^. Further work should focus on determining whether variability in IFN-I and *MAVS* signaling among individuals can explain the differential susceptibilities to CHIKV-induced cardiac complications.

Multiple PRRs are involved in the *in vivo* control of CHIKV infection^30^. Here, we demonstrate that CHIKV clearance from infected tissue is MAVS-dependent, but independent of MYD88, TRIF, or the NLRP3 inflammasome. However, we found reduced number of infectious particles isolated from infected hearts of Casp1*/11*^*-/-*^ and *Myd88*^*-/-*^ mice compared to WT, suggesting that the NLRP3 inflammasome as well as signaling through TLR7 may play a role in the early steps of CHIKV-heart infection. We demonstrate that in the absence of MAVS, CHIKV persist in the cardiac tissue. Intriguing, systemic levels of both IFNα and IFNβ were significantly higher in *Mavs*^*-/-*^ mice compared to WT, and directly correlated with the viral loads observed in *Mavs*^*-/-*^. Multiple PRRs are involved in the IFN-I production and protection associated with alphavirus infection^31,49^, suggesting that other PRRs, such as TLR3 or TLR7 may be contributing to the rescue in the production of systemic IFN-I in this model^50^. In line with this, we found that in *Mavs*^*-/-*^ mice, infected hearts display similar levels of local *Isg15* compared to WT, indicating that cardiac tissue is responding to IFN-I in the absence of MAVS signaling. These data suggest that either: (i) the delay in CHIKV clearance from cardiac tissue does not entirely depend on the reduced IFN-I induction associated with *Mavs*^*-/-*^ cells, or (ii) the bulk nature of our study does not provide the resolution to define local effects associated to *Mavs*^*-/-*^ specific responses. An example of the latter was observed for Rift Valley fever virus-infected brains, in which no differences in the global induction of the IFN-I signaling between *Mavs*^*-/-*^ and WT were observed by bulk RNA-seq^51^, but became apparent when analyzed at single-cell resolution^51^. Single-cell technologies have been recently applied to the study of cardiac tissue and viral myocarditis^42^. Thus, studies on CHIKV-infected cardiac tissue at single-cell level may help elucidate the role of the local cellular effects in CHIKV clearance. *Mavs*^*-/*-^ mice display different susceptibilities to related arthritogenic alphaviruses^30,52^. While *Mavs*^*-/-*^ mice infected with Ross River Virus developed severe disease and succumbed to infection^52^, CHIKV infected mice survived when inoculated subcutaneously^30^ (**Extended Data Table 5**). Comparative studies addressing the role of PRR in controlling the infection of related arthritogenic alphavirus would be fundamental for expanding our knowledge virus-specific mechanisms of pathogenesis.

We demonstrated that CHIKV infection led to focal and multifocal myocarditis in *Mavs*^*-/-*^ mice. Yet, serum levels of cTnT remained unaltered, supporting the absence of cardiac myocyte damage. We found that CHIKV does not infect cardiac myocytes in mice, consistent with the lack of evidence of cardiac myocyte infection in human post-mortem cardiac tissue samples^12^. However, elevated cardiac troponins have been observed in individuals with CHIKV-associated cardiac manifestations^6,7,13,16^, raising the possibility that under certain conditions CHIKV can induce cardiac myocyte damage. Another alternative is that human and murine cardiac myocytes have different susceptibilities to CHIKV infection. Further studies using human heart tissue explants or human cardiomyocyte organoids are required to evaluate the relative susceptibilities of human cardiac myocytes to CHIKV infection.

Finally, we found that CHIKV infection led to chronic vascular inflammation in major vessels attached to the base of the heart. This chronic inflammation in the vasculature is intriguing and highlights the vasculature itself as a potential niche for CHIKV infection. Indeed, we detected a differential behavior in CHIKV susceptibilities between the top and the bottom sections of the heart, and CHIKV active replication was detected in cells within vessels up to 10 dpi in *Mavs*^*-/-*^. These results demonstrate that persistent infection in cardiac tissue leads to cardiac inflammation and potentially cardiac tissue damage, and highlights a potential risk to develop long-term vascular damage associated to CHIKV infection.

Overall, this study demonstrates the direct causality between CHIKV active replication in the heart and the development of cardiovascular manifestations. Future studies focalized on atypical manifestations of arboviral diseases and their pathogenesis are fundamental for understanding the full spectrum of the consequences of these infections. This knowledge is paramount for monitoring of atypical manifestations in endemic areas as well as in future epidemics. In turn, this could be leveraged for early diagnostics, prevention, and therapeutic interventions especially in endemic areas where the circulation of these viruses represents a public health burden. Altogether, here we provide mechanistic insight into how CHIKV can lead to cardiac damage, underscoring the importance of monitoring cardiac function in patients with CHIKV infections and laying the foundation for the development of new approaches to prevent viral-induced cardiac complications.

## ACKNOWLEDGMENTS

We would like to thank all members of the Stapleford lab for helpful comments on this manuscript. We thank NYUMC Experimental Pathology Research Laboratory, Mark Alu, and Branka Brukner Dabovic. We are also grateful to the NYU Genome Technology Center for assistance during sequencing and the NYU Langone Microscopy Laboratory for assistance during imaging. We thank Drs. Meike Dittmann and Ken Cadwell who kindly provided essential equipment and reagents. We thank Drs. Dominick Papandrea and Ludovic Desvignes for BSL3 assistance. We thank Drs. Ken Cadwell and Victor Torres for critical reading of the manuscript. This work was supported by a start-up package from the NYU Grossman School of Medicine (K.A.S), NIH/NIAID R01 AI162774-01A1 (K.A.S) and 1R01AI143861 (K.M.K), the NYU Cardiovascular Research Center Program pilot grant (K.A.S. and M.G.N), the American Heart Association Postdoctoral Fellowship 19-A0-00-1003686 (M.G.N.), Public Health Service Institutional Research Training Award T32 AI 7180-39 (B.A.R-R.), Training Program in Immunology and Inflammation Training Award 5-T32 AI100853-10 (E.B.). The Experimental Pathology Research Laboratory, the NYU Langone Microscopy Laboratory and the NYU Genome Technology Core are partially supported by NYU Cancer Center support grant P30CA016087 and by NYU Langone’s Laura and Isaac Perlmutter Cancer Center.

## AUTHOR CONTRIBUTIONS

M.G.N. and K.A.S conceived the project and designed experiments; M.G.N, S.N.S, E.B, S.T.Y, P.D.Y, B.A.R.R. performed experiments, F.I. analyzed RNA-seq data; M.G.N, E.B., F.I., and K.A.S. analyzed data, C.L. oversaw histology experiments, N.N. performed histopathology analysis; V.M. develop the Visiopharm app for histology analysis; M.Z.D. performed slide cryosections; K.K and K.A.S supervised the project. M.G.N. and K.A.S. wrote the original draft, and all authors were involved in manuscript review and editing.

## DATA AND CODE AVAILABILITY

Bulk RNA sequencing data sets generated in this study are available at GEO: GSE204689. Code is available at GitHub: https://github.com/fizzo13/CHIKV. Any additional information required to reanalyze the data reported in this paper is available from the lead contact upon request.

## MATERIALS AND METHODS

### Biosafety

Work with CHIKV was performed in a biosafety level 3 (BSL-3) laboratory at NYU Grossman School of Medicine.

### Cells

Cells were maintained at 37ºC in 5% CO_2_. Baby hamster kidney (BHK-21, ATCC CCL-10) were grown in Dulbecco’s modified Eagle’s medium (DMEM; Corning) supplemented with 10% fetal bovine serum (FBS; Atlanta Biologicals) and 1% nonessential amino acids (NEAA, Gibco). Vero cells (ATCC CCL-81) were grown in DMEM with 10% newborn calf serum (NBCS; Gibco). Human primary cardiac cells were obtained from PromoCell and grown following the company recommendations. Human Cardiac Fibroblast (C-12375), human Cardiac Microvascular Endothelial Cells (C-12285), and human Cardiac Myocytes (C-12810) were grown in Fibroblast Growth Medium 3 (PromoCell, C-23025), Endothelial Cell Growth Media MV (PromoCell, C22020), and Myocyte Growth Media (PromoCell, C22070), respectively. All the cell types were confirmed free of mycoplasma using Lookout Mycoplasma PCR detection kit (Sigma-Aldrich).

### Viruses

CHIKV (CHIKV strain 06-049; AM258994) and CHIKV reporter (CHIKV-ZsGreen; CHIKV strain 06-049) infectious clones were generated as previously described^20,53^. *Virus rescue from infectious clones*. Briefly, 10 μg of each of the infection clone plasmids were linearized overnight with NotI (Thermo-Scientific) at 37ºC, phenol:chloroform extracted, ethanol precipitated, and resuspended in nuclease-free water at 1 μg/μl. *In vitro* transcribed viral RNAs were produced using the SP6 mMESSAGE mMACHINE kit (Ambion) following the manufacturer’s instructions. After RNA synthesis, samples were DNAse treated, and RNAs were purified by phenol:chloroform extraction, ethanol precipitation, and resuspended in nuclease-free water at a concentration of 1 μg/μl. RNA integrity was confirmed by gel electrophoresis (1% agarose). RNA was aliquoted and stored at -80 ºC until electroporation. Infectious virus was produced by electroporating BHK cells with *in vitro* transcribed viral RNAs. For that purpose, cells were trypsinized, washed twice with ice-cold DPBS (Corning), and resuspended at 1×10^7^ cells/ml in PBS. 390 μl of cells were mixed with 10 μg of *in vitro* transcribed RNA and added to a 2 mm electroporation cuvette (BioRad). BHK-21 cells were electroporated with 1 pulse at 1.2 kV, 25 μF, with infinite resistance. Cells were allowed to recover for 10 min at room temperature (RT), transferred into 6 ml of warm DMEM supplemented with 10% FBS and 1% NEAA, and placed in a T25 flask at 37ºC for 72 h. Virus was harvested, clarified at 1,200 x g for 5 min, and 3 ml of the clarified supernatants were used to infect T175 flask to generate working stocks (Passage 1). Viral titers were determined by plaque assay in Vero cells.

### Plaque assay

10-fold dilutions of each virus in DMEM were added to a monolayer of Vero cells for 1 h at 37ºC. Following incubation, cells were overlaid with 0.8% agarose in DMEM containing 2% NCBS and incubated at 37ºC for 72 h. The cells were fixed with 4% formalin, the agarose plug removed, and plaques visualized by crystal violet staining. Titers were determined by counting plaques on the highest countable dilution. To generate heat-inactivated virus (HI), virus stocks with titers of 10^6^ PFU/ml were incubated in a water bath at 56ºC for 4 h. Loss of infectivity was confirmed by plaque assay on Vero cells.

### *In vitro* IFN-I response experiments

Human primary cardiac cells were seeded into 96-well flat transparent black plates (Corning, 07200588) at a density of 18,000 cells/well. After 24 h cells were treated as follows: (i) *In vitro* poly (I:C) stimulation, (ii) Ruxolitinib treatment, or (iii) mock treatment. For poly(I:C) stimulation, human cardiac fibroblasts in 96-well plate were transfected with 62 ng of high molecular weight poly(I:C) (InvivoGen) using TransIT-mRNA Transfection Kit (Mirus Bio), or treated with TransIT transfection alone (carrier control). After 2h cells were washed and used for downstream experiments. For Ruxolitinib condition, cells were pre-treated with 5 μM Ruxolitinib (Invitrogen) or mock-treated for 3 h, cells were washed and used for downstream experiments. Cells pretreated with poly(I:C), Ruxolitinib or mock-treated were then infected with CHIKV-ZsGreen at an MOI of 0.25 or mock-infected with DMEM for 1 h at 37 ºC. After 1h, the inoculum was removed, monolayers were washed once with cold PBS, and 100 μl of Fibroblast Growth Medium 3 (PromoCell, C-23025) was added to the cells. Fibroblast Growth Medium 3 with 5 μM Ruxolitinib was added to the cells for the Ruxolitinib condition. Cells were incubated at 37 ºC with 5 % CO2 for 24 h. Cells were fixed with 10% formalin for 1 h, and stained for microscopy (see below).

### *In vitro* recombinant IFN-I pre-treatment experiments

Human primary cardiac cells were seeded into 96-well plates flat transparent black plates (Corning, 07200588) at a density of 18,000 cells/well. After 24 hs cells were stimulated with the indicated concentrations of IFNα A/D (Millipore Sigma, #I4401) or IFNβ (Millipore Sigma, #IF014) for 24 h. Cells pretreated with IFN or mock-treated were infected with CHIKV-ZsGreen at an MOI of 0.25 or mock-infected with DMEM for 1 h at 37 ºC. After 1h, the inoculum was removed, monolayers were washed once with cold PBS, and 100 μl of Fibroblast Growth Medium 3 (PromoCell, C-23025) was added to the cells. Cells were incubated at 37 ºC with 5 % CO2 for 24 h. Cells were fixed with 10% formalin for 1 h, and stained for microscopy (see below).

### Staining and high content microscopy analysis

After fixation, cells were washed two times with PBS, permeabilized with 0.1% Triton-X/PBS for 10 min, and washed again with PBS. For recombinant IFN-I pretreatment experiments, cells were stained with DAPI (Thermo) at a dilution of 1:1000 for 10 min at room temperature and washed three times with PBS. For *in vitro* response experiments, cells were blocked overnight in 1%BSA/PBS at 4 ºC. Cells were stained with anti-ISG15 antibody (ProteinTech, 15981-1-AP) at a dilution of 1:100 in 0.1%BSA/PBS for 1 h at 37 ºC, and washed three times with PBS. Cells were stained with secondary AF555-anti-Rabbit antibody (Invitrogen) and DAPI (Thermo) at 1:1000 dilution for 1 h at 37 ºC and washed three times with PBS. Plates were imaged using the CellInsight CX7 High-Content Screening Platform (Thermofisher) with the 4x objective in 9 fields covering completely each well from the 96-well plate. The high-content software was used for microscopy and imaging analysis.

### Mosquitoes

*Aedes (Ae*.*) albopictus* eggs from Tallman Island Wastewater Treatment Plant in Queens, New York City, were collected by the New York Department of Health^54^. Mosquitoes were reared and maintain in Memmert humidified chambers at 28 ºC with 70% humidity and 12 h diurnal light cycle. All the transmission studies were done with mosquitoes in generation F7. *Mosquito infection. Ae. albopictus* mosquitoes from Tallman (F7) were exposed to an artificial blood meal containing 10^6^ PFU/ml CHIKV. Briefly, viruses were mixed 1:2 with PBS-washed sheep blood (Fisher Scientific, MA, USA) supplemented with 5 mM ATP. Mosquitoes exposed to non-infectious blood meals were used as non-infected controls. Female mosquitoes were allowed to feed on 37 ºC blood meals through an artificial membrane for 60 to 90 min. Engorged females were identified, sorted and incubated at 28 ºC with 10% sucrose ad libitum for 7 days. Prior to transmission, mosquitoes were food deprived for 12 h.

### Mice

All mice were housed with food and water ad libitum under 12 h dark/light cycle in a pathogen-free facility at NYU Grossman School of Medicine. Experimental animals were housed in groups of up to 5 mice per cage. Mice were bred and grown in house and maintained in the same facility under specific pathogen-free conditions included the following strains: C57BL/6J wild type mice (000664, Jackson Laboratories). C57BL/6(027) wild type mice (Charles River Laboratories), *Ifnar1*^*-/-*^ (028288, Jackson Laboratories); *Myd88*^-/-^ (009088, Jackson Laboratories); *Casp1/11*^*-/-*^ (016621, Jackson Laboratories); *Mavs*^*-/-*^ (008634, Jackson Laboratories), *Trif*^*Lps2/Lps2*^ (005037, Jackson Laboratories). All mice were bred as heterozygotes to generate WT and KO littermate controls, and subsequent WT or KO colonies coming from littermate controls were used for experiments. *Mouse infections*. Animal experiments were performed in accordance with all NYU Grossman School of Medicine Institutional Animal Care and Use Committee guidelines (IACUC) and in accordance with guidelines from the National Institutes of Health, the Animal Welfare Act, and the U.S. Federal Law. All mouse studies were performed under BSL-3 conditions. Infections were carried out by either subcutaneous inoculation, mosquito bite, or intravenous injection via tail vein injections. Briefly, 6-8-week-old male and female were infected with 10^5^ PFU of CHIKV, CHIKV-ZsGreen, or mock-infected with carrier (PBS or DMEM) or similar inoculum of heat inactivated virus (HI). For subcutaneous inoculation, mice were anesthetized by inhalation of isoflurane (Henry Schein Animal Health), and injected with 50 μl of virus or control in the left rear footpad. For intravenous inoculation mice were restrained and injected via tail vein injection with 100 to 200 μl of virus or control inoculum using a 29G insulin syringe (Fisher Scientific). For natural transmission, mice were immobilized over a mesh covered pint cup containing 4 to 8 CHIKV-infected or uninfected (blood only) mosquitoes, and mosquitoes were allowed to feed for 40 min. Afterwards, mice were returned to their cages, mosquitoes were killed and homogenized, and viral titers of the infected mosquitoes were determined by plaque assay. Infected experimental mice were euthanized at different timepoints post infection by CO_2_ inhalation. Blood was collected by cardiac puncture, and mice were immediately dissected for transcardiac perfusion. Mice were perfused with 30 ml of cold PBS with a 21G to 23G needle. Quality of cardiac perfusion was assessed by confirming full blood removal in liver and kidneys. Then, hearts were extracted and placed in a 12-well plate with 1 ml PBS. Each heart was opened in four chambers and confirmed free of any blood clot. All the mice in this study were perfused with the exception of those used for histopathology and immunostaining. Heart, calf muscle and pancreas were collected in 500 μl of PBS containing one (heart and pancreas) or two (calf muscle) 5 mm stainless steel beads (QIAGEN), homogenized with TissueLyser II (QIAGEN) at 30 1/s for 2 min (heart and pancreas) or 4 min (calf muscle). Debris was removed by centrifugation at 8,000 rpm for 8 min. Blood was collected in 1.5 ml tubes containing 20 ml of 0.5 M EDTA and centrifuged at 8,000 rpm for 8 min. Viral titers were quantified by plaque assay in Vero cells or TCID_50_ in BHK-21 cells.

### Median Tissue Culture Infectious Dose (TCID_50_)

Briefly for TCID_50_ determination, BHK-21 cells were seeded into 96-well plates (10,000 cells/well). For heart, muscle and pancreas samples the initial dilution corresponds to a dilution 1:5 from the original homogenate, for serum the initial dilution was 1:20. After the first dilution, ten-fold dilutions of tissue homogenates or serum were made in DMEM supplemented with 1% Antibiotic-Antimycotic (Anti-Anti, Invitrogen). Monolayers were infected with 50 μl of the dilutions. After 1 h, 100 ml of DMEM + 10% FBS + 1% Anti-Anti was added to each well. Cells were incubated at 37 ºC with 5 % CO2 for 96 h. Plates were inspected under the microscope for evidences of cytopathic effect (CPE). Non-infected tissue was used as negative control, and mock treated wells were included in each plate. Positive wells, corresponds to any well showing CHIKV specific CPE. Negative wells, corresponds to any well with no CPE. Following incubation, cells were fixed with 10% formalin, and visualized under the microscope. Each sample was measured in 6 or 7 technical replicates and the calculation was performed using the online TCID_50_ calculator (https://www.klinikum.uni-heidelberg.de).

### RNA extractions and RT-qPCR

RNA extractions were performed using TRIzol™ reagent (Invitrogen, 200 μl of clarified tissue homogenate was added to 500 μl of TRIzol™) following the manufacturer’s guidelines. Extracted RNA was quantified and diluted to 200 ng/μl. For Taqman assays, a standard curve spanning a range of 100 ng/μl to 1E-6 ng/μl of CHIKV viral RNA was generated for each dataset using *in vitro* transcribed CHIKV RNAs as described above. For the different tissues, we optimized the amount of total viral RNA to be within the linear range of the calibration curve. The number of genomic (nsp4) or subgenomic (E1) CHIKV RNA was quantified by RT-qPCR using 0.25-1 μg of total RNA as template and the Taqman RNA-to-CT One-Step kit (Applied Biosystems™, Beverly, MA, USA) in a total reaction volume of 25 μl/well. The following primers and probes were used: CHIKV primers to amplify nsP4 fragment (CHIKV forward (IOLqPCR6856F): 5’-TCACTCCCTGCTGGACTTGATAGA-3’; reverse (IOLqPCR6981R): 5’-TTGACGAACAGAGTTAGGAACATACC-3’) and aprobe (5′-(6-carboxyfluorescein)-AGGTACGCGCTTCAAGTTCGGCG-(black-holequencher)-3′). CHIKV primers to amplify E1 fragment (CHIKV forward (IOLqPCR10865F): 5′-TCGACGCGCCCT CTTTAA-3′); reverse (IOLqPCR10991R): 5′-ATCGAATGCACCGCACACT-3′ and a probe 5′-/56-FAM/ACCAGCCTG/ ZEN/CACCCATTCCTCAGAC/ 3IABkFQ/-3′). The final concentration of primer and probes were 1 μM and 0.6 μM, respectively. Reverse transcription was carried out on QuantStudio 3 qPCR instrument using the following protocol: 48 ºC for 30 min, followed by 10 min at 95 ºC. Amplification was accomplished over 40 cycles as follows: 95 ºC for 15 sec, 60 ºC for 1 min. For Sybr green quantifications, levels of *Isg15, Mx1, ApoD, Ccl5, and Ifitm3* in heart homogenates were measured by SYBR Green Master Mix (Thermo) and following manufacturer’s instructions on a QuantStudio 3 qPCR instrument. Briefly, cDNA synthesis was performed on purified RNA samples as described above using the Maxima H minus-strand kit (Thermo Scientific) and random primers, using the following protocol: 25 ºC for 10 min, 50 ºC for 30 min, and 85 ºC for 5 min. Amplification was accomplished over 40 cycles as follows: 95 ºC for 15 sec, 60 ºC for 1 min. The melt curve was evaluated for each reaction. Primers: *Isg15* (fw:5’-GGTGTCCGTGACTAACTCCAT-3’, and rv:5’-TGGAAAGGGTAAGA CCGTCCT-3’), *Mx1* (fw:5’-TCCCTGGGAAGGAGTGAAGCCAG-3’, and rv:5’-CTTGCCACTGGGGTCAGGCAC-3’), *ApoD* (fw:5’-CTGGGTGGAAACTTCAGTCATCTGATCT-3’, and rv: 5’-CGCCAGCGTGGCCAGGAAC-3’), *Ccl5* (fw:5’-TGCCCACGTCAAGGAGTATTTC-3’, and rv:5’-TCCTAGCTCATCTCCAAATAGTTGATG-3’), *Ifitm3* (fw:5’-TTCAGTGCTGCCTTTGCTC-3’, and rv:5’-CCTTGATTCTTTCGTAGTTTGGGG-3’) and *18S* (fw:5’-GTAACCCGTTGAACCCCATT-3’, and rv: 5’-CCATCCAA TCGGTAGTAGCG-3’). The relative expression of the respective genes to 18S expression was calculated using the ΔΔCT method, and values were expressed as fold change normalized to mock-infected control. ΔΔCT for each sample (infected and mock-infected) was calculated against the average of all the ΔCT values calculated for the mock condition. **RNA extractions and RNA-sequencing analysis**. For bulk RNA-sequencing C57BL/6 male mice were infected intravenously with 10^5^ PFU of CHIKV or HI control, and hearts were perfused with 30 ml of PBS and collected at 2 and 5 dpi. Hearts were collected in 500 μl of TRIzol™ (Invitrogen) with one 5 mm stainless steel bead and processed as described above. RNA was extracted using Zymo direct-zol RNA purification kit (Zymo research, #R2072) according to the manufacturer’s instructions. A total of 500 ng per sample was sent to the NYU Genome Technology Center for quality control assessment, automated TruSeq stranded total RNA, with RiboZero Gold library preparation, and Illumina Novaseq6000 sequencing (single pair 100). FASTQ files were aligned to a concatenated file containing the mouse genome (version mm10) and the CHIKV genome (AM258994) through STAR (v2.5.2) in two pass basic mode using the gencode transcript annotation (vM25). Structural and non-structural CHIKV polyprotein coordinates (CHIKVgp1 and CHIKVgp2) were added to the transcriptome annotation, and the transcript count matrix was generated using the summarize overlaps function form the Genomic Alignments (v1.22.1) package. Differential gene expression analysis based on negative binomial distribution was performed using DEseq2 (v1.26.0). For pathways enrichment analysis, genes were ranked based on -Log_10_(FDR) values, assigning a positive or negative value depending on the direction of change. The pathway lists were obtained from the msigdbr (v7.2.1) package (Hallmark, REACTOME or KEGG) and used as input for GSEA pathway enrichment analysis through the fgsea (v1.12.0) R package. Volcano plots were generated using ggplot2 (v3.3.3) and heatmaps were generated using pheatmap (v1.0.12).

### Tissue preparation for immunofluorescence and fluorescence microscopy

Hearts were removed from euthanized mice and fixed in PFA, lysine and periodate buffer (PLP, 0.05 M phosphate buffer, 0.1 M L-lysine, pH 7.4, 2 mg/ml NaIO_4_, and 10 mg/ml paraformaldehyde) at 4°C for 24 h. Next, tissues were dehydrated in 30% sucrose overnight at 4°C and subsequently embedded in OCT media. Frozen tissue sections were sectioned at 5 μm thickness at the NYU Experimental Pathology Research Laboratory. Fc receptors were blocked with 0.5% anti-CD16/32 Fc block antibody (Biolegend, clone 93) diluted in PBS containing 2% goat serum (Vector Laboratories, S-1000), 2% FBS (Atlanta) and 0.25% Triton-X 100 (Fisher, BP151-100) for 1 h at RT. Sections were washed 5 times with the same buffer, and treated with Trueblack Lipofuscin autofluorescence quencher (Biotium, CAT23007) in ethanol 70% (5% v/v) for 1 min. Sections were washed 5 times and stained with eF570-αSMA (Invitrogen, clone: 1A4) with either APC-CD45 (BioLegends, clone: 30-F11), AF647-TNT (BD Pharmingen, clone: 13-11), AF647-CD31 (BioLegends, clone: MEC13.3) or AF647-Vimentin (Abcam, clone: EPR3776) for 1 h, at RT. All antibodies were used at 1:100 dilution. For IFITM3 or ISG15 staining, slides were stained with 1:100 anti-Fragilis antibody (Abcam, ab15592, polyclonal) or 1:100 anti-ISG15 antibody (Invitrogen, PA5-79523, polyclonal) for 1h at RT, respectively; washed and stained with a secondary AF555-anti-Rabbit antibody (Invitrogen) at 1:1000 dilution for 1 h. All the slides were then stained with DAPI (Thermo) at 1:1000 dilution. Sections were washed with the same buffer and mount for microscopy analysis. Images were acquired using a Zeiss LSM 880 confocal microscope (Carl Zeiss).

### Imaris surface colocalization analysis

The imaging data were processed and analyzed using Imaris software version 9.8 (Bitplane; Oxford Instruments). Briefly, surfaces were built for each channel using specific threshold, number of voxels and surface grain size. The selection of parameters was dependent on each experimental set of images. Once the parameters were defined batch analysis was performed to calculate the total amount of overlapping surfaces between two channels. Analysis was performed on 2×2 tiled images acquired at 20x magnification from the Zeiss 880 confocal microscope, where 2 to 4 different sections per slide were imaged from two to three independent animals per condition.

### Histopathology

The heart and part of the ascending thoracic aorta were removed from euthanized mice, fixed in 10% buffered formalin (Fisher Scientific) for 72 h, and processed through graded ethanol, xylene and into paraffin in a Leica Peloris automated processor. Five-micron paraffin-embedded sections were cut parallel to the long axis of the heart from two to three distinct levels (400 μm, 800 μm, and 1200 μm; or 800 μm, and 1200 μm). Sections of the 4-chamber heart were deparaffinized and stained with hematoxylin and eosin (H&E) on a Leica ST5020 automated histochemical stainer or immunostained on a Leica BondRX® autostainer, according to the manufacturers’ instructions. In brief, sections for immunostaining underwent epitope retrieval for 20 min at 100 ºC with Leica Biosystems ER2 solution (pH9, AR9640) followed by a 30 min incubation at 22 ºC with either anti-CD11b (Novus, NB110-89474) diluted 1:10,000, with anti-CD3 (CST, 78588S; clone E4T1B) or with anti-cleaved CASP3 (CST, 9579S, clone D3E9) diluted 1:1,000. The primary antibodies were detected using the BOND Polymer Refine Detection System (Leica, DS9800). As positive controls of the IHC, sections of mice spleen were stained with anti-CASP3, anti-CD3 and anti-CD11b. Sections were counterstained with hematoxylin scanned on either a Leica AT2 or Hamamatsu Nanozoomer HT whole slide scanners and imported into the NYU Omero image database for viewing and annotation. The severity of tissue pathology and determination of the different categories described in Table S1 and S5, were blindly analyzed by a pathologist (N.N.). Masson trichrome staining was performed as previously described^55^. In brief, deparaffinized slides were further fixed in Bouin solution for 1 h at 60 ºC. The slides were then serially stained with Weigert Hematoxylin (Polysciences, 25088B1&B2) for 10 min, followed by Biebrich Scarlet-Acid Fuchsin (Polysciences, 25088C) for 1 min and Phosphotungstic acid (Polysciences, 25088D) for 10 mins with washes between each step. Slides were then transferred directly into aniline blue (Polysciences, 25088E) for 5 min and differentiated in 1% acetic acid (Polysciences, 25088f) for 1 min before dehydration and mounting with permount (Fisher, SP15-100).

### CASP3 IHC quantification analysis

To determine the number of cleaved CASP3^+^ cells whole slide scans were analyzed using Visiopharm^R^ image analysis software version 2021.09. Images were imported to Visiopharm^R^’s database. Manual outlining of regions of interest (ROI) was performed to divide the four-chamber heart section into right, left ventricles, septum and atria. A Visiopharm^R^ app was generated to detect DAB-stained cells in the mouse heart. Briefly, image pre-processing steps were run to create features enhancing DAB signal and nuclear signal through color deconvolution and filtering. Segmentation was achieved using a trainable linear bayesian algorithm. Post-processing steps were applied to define nuclei as positive or negative. Results were provided as number of total nuclei, number of positive and number of negative nuclei per ROI analyzed.

### Cytokine and chemokine analysis

Mice were infected intravenously with 10^5^ PFU of CHIKV or HI control, and heart were collected at 2 and 5 dpi. Heart tissue homogenates generated as described above were diluted 1:5 in 5X PBS:HALT protease inhibitor EDTA-free (Thermo). Total amount of protein was quantified using BCA protein assay (Thermo Fisher Scientific) and samples were diluted to 1.5 mg/ml in PBS:HALT protease inhibitor, 75 ml aliquots were made and stored at -80ºC until needed. Heart homogenates were analyzed for cytokine and chemokine levels using a Bio-Plex Pro Mouse Cytokine 23-plex Assay kit (Bio-Rad, #m60009rdpd) and ProcartaPlex mouse IFN-α/IFN-β 2-plex (ThermoFisher, EPX02A-22187-901) according to the manufacturer’s instructions.

### Mouse Cardiac troponin-T ELISA

Mice were infected intravenously with 10^5^ PFU of CHIKV or HI control, and blood collected as described above, incubated at RT for 30 min, and centrifuged at 1000 x g for 15 min at 4ºC. The upper phased was then transferred to a clean tube and centrifuged at 10,000 x g for 10 min. Samples were moved to a clean tube and then 50 ml aliquots were made and stored at -80ºC until needed. Serum was analyzed for mouse cardiac troponin-T using a precoated microtiter plate (amsbio, AMS.E03T0017) and using a recombinant cardiac troponin-T standard curve according to the manufacturer’s instructions.

### Single-cell analysis of *Tabula Muris* datasets

Data from *Tabula muris* (Tabula Muris et al., 2018) was downloaded from https://tabula-muris.ds.czbiohub.org. Data was analyzed using the Seurat (v3.2.1) R package. Briefly, data was normalized using the “Normalize Data” function with default parameters. After normalization, we applied the function “Find Variable Features” to identify genes of interest used for downstream dimensionality reduction via the function “RunPCA” and clustering through the “Find Neighbors” and “Find Clusters” functions, with default parameters. The original cluster annotations from *Tabula Muris* were maintained. Dot plot for normalized expression of potential cell markers was generated using the “DotPlot” function.

### Statistical analysis

Statistical significance was assigned when *P* values were < 0.05 using Prism Version 9 (GraphPad) or R-studio (v3.6.0). Specific tests with exact *P* values are indicated in the Figure legends. All experiments were performed at least in biological duplicates.

## EXTENDED FIGURES

**Extended Data Fig. 1.**
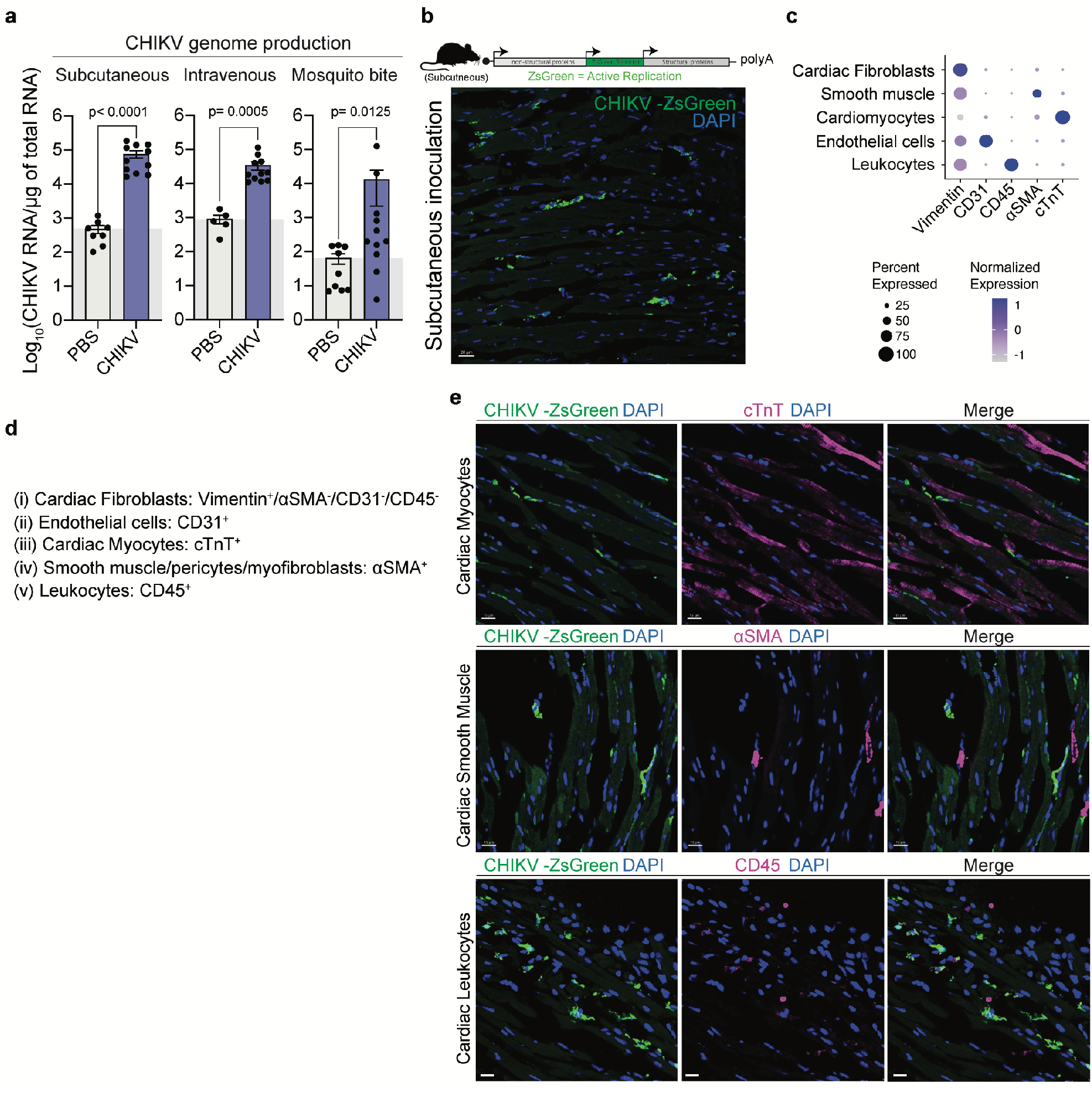
a. (Related to Fig. 1). **a**. Six-week-old C57BL/6 mice were inoculated with 1E5 PFU of CHIKV or mock-infected (PBS) subcutaneously, via natural transmission through a mosquito bite, or intravenously. Mice were harvested at 2 dpi for subcutaneous and intravenous infection routes, and at 2 and 3 dpi for mice infected through mosquito bite. Animals were perfused with 30 ml of cold PBS. Viral RNA genomes in heart tissue were quantified using RT-qPCR. Data represent two to three independent repetitions. N=5-11 mice/group. The gray box represents the background RNA levels determined for mock-infected controls. **b**. Upper panel: Schematic representation of the experimental setup. Mice were inoculated subcutaneously with 1E5 PFU of reporter CHIKV-ZsGreen or mock (PBS). Bottom panel: Examples of infected myocardium. Blue channel: DAPI. White arrows: CHIKV-infected cells. Scale= 20 μm. **c**. Dot plot showing the expression levels of the markers used in the study by cardiac cell type using the *Tabula Muris* data set. **d**. Description of cellular markers used to define each cardiac cell type. **e**. Representative fluorescence microscopy images of myocardium of CHIKV-infected hearts stained with cardiac myocytes markers (top panel), cardiac smooth muscle cells (middle panel), or cardiac leukocyte cells markers (bottom panel). The panels represent the individual or merge channels. Scale= 15 μm.

**Extended Data Fig. 2.**
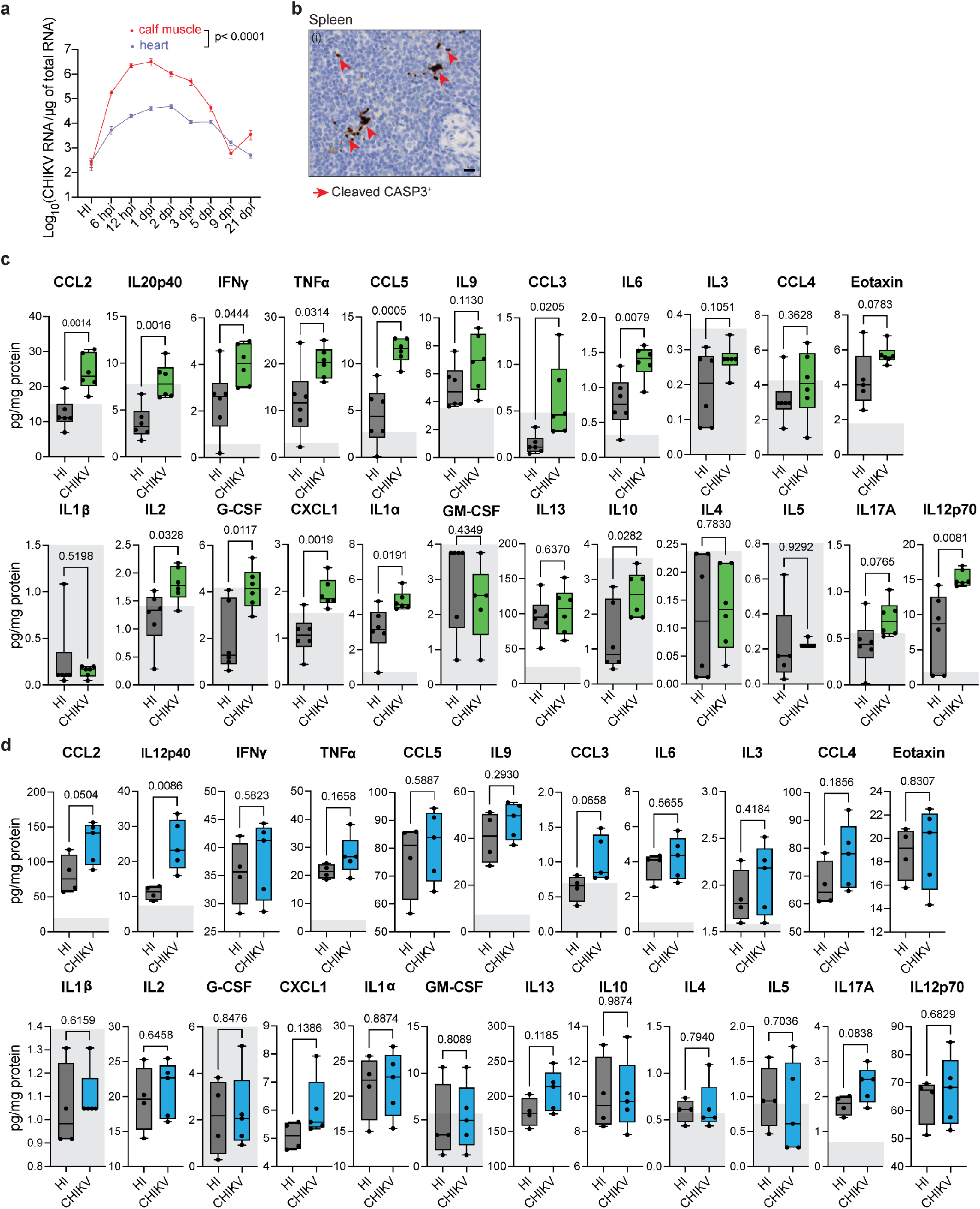
(Related to Fig. 2). **a**. CHIKV replication kinetics in the heart (blue) and the calf muscle (red) measured by RT-qPCR. Each time point represents the average and the standard deviation of the mean. The gray box represents the background RNA levels determined for HI controls. N=5-10 mice/group. **b**. Positive control of cleaved CASP3 antibody staining used for IHC. **c-d**. Individual values from which **Fig.2d** was generated. Each bar graph represents the protein levels in cardiac tissue homogenates for different analytes from the Bio-Plex Pro Mouse Cytokine 23-plex Assay. Values within the gray shading represent concentrations of the analyte that were beyond the calibration curve and therefore determined by extrapolation of the curve by the Luminex Software. Mock: HI inoculum. N=5-6 mice/group. P values were calculated using mixed-effects model test **(a)** or using Mann-Whitney test **(c-d)**.

**Extended Data Fig. 3.**
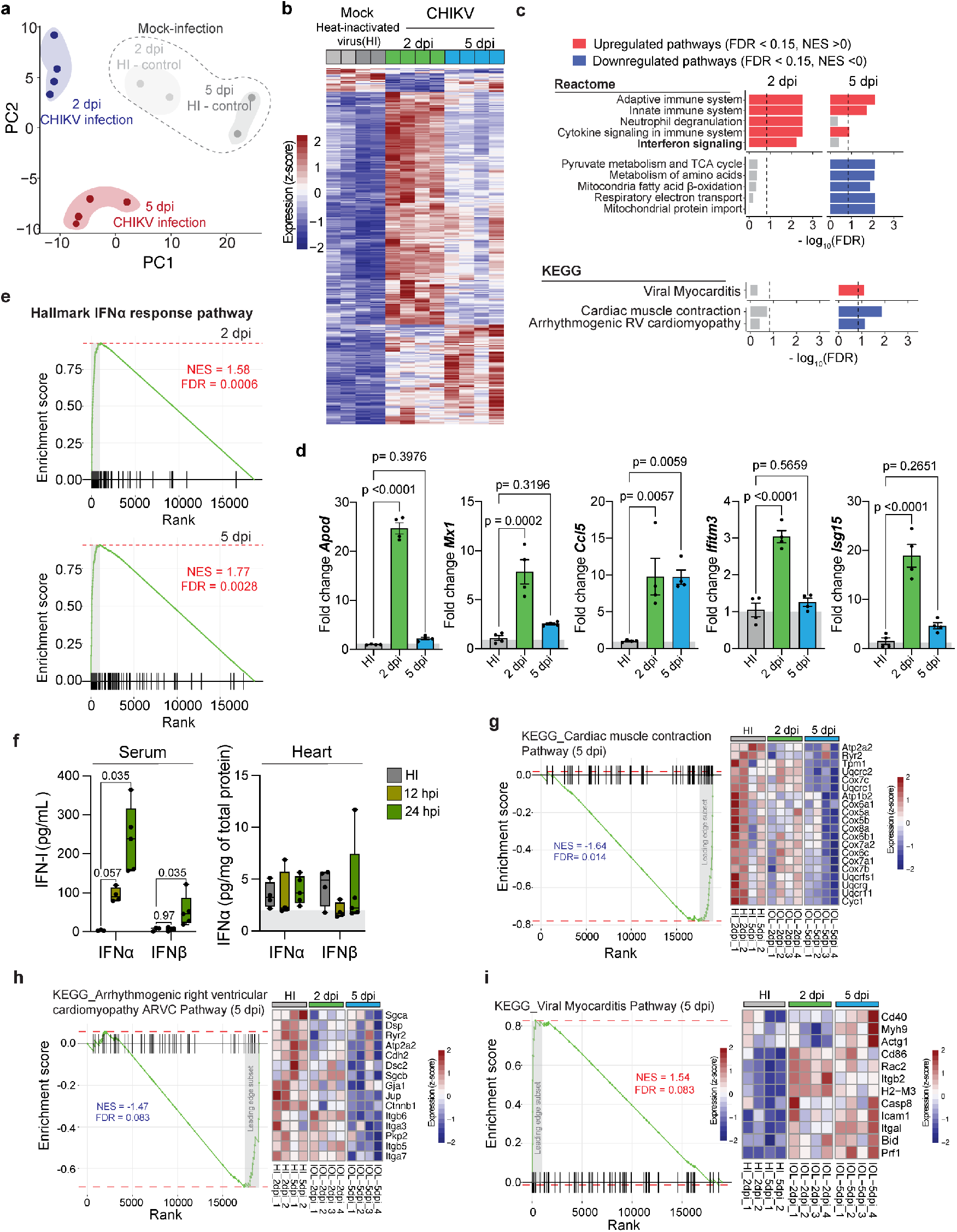
(Related to Fig. 2 and Fig.3). **a**. Principal component analysis (PCA) showing that biological replicates clustered by treatment. **b**. Heatmap showing differentially expressed genes (FDR < 0.15 and absolute log_2_ FC > 1). The boxes on top indicate the biological replicates for mock (heat-inactivated virus; N=4; grey boxes) and CHIKV-infected at either 2 dpi (N=4; green boxes) or 5 dpi (N=4; blue boxes). **c**. GSEA pathway enrichment analysis for Reactome, and KEGG datasets showing top and downregulated pathways at 2 dpi and 5 dpi. **d**. Validation of RNA-seq targets. *Apod, Mx1, Ccl5, Ifitim3*, and *Isg15* expression levels from CHIKV-infected and mock-infected (HI inoculum) heart homogenates were measured by RT-qPCR at 2 and 5 dpi. Gray boxes indicate fold change of 1. N=4 mice/group. **e**. Enrichment plot for the IFNα response pathway (Hallmark) at 2 dpi (upper panel) and 5 dpi (bottom panel). **f**. Protein levels of IFN-α and INF-β in serum (left panel) and heart tissue homogenates (right panel) at 2 dpi. Values within the gray shading represent concentrations of the analyte that were beyond the calibration curve and therefore determined by extrapolation of the curve by the Luminex Software. Dataset at 24 hpi are the same as the dataset for WT at 24 hpi in **Fig.4h-i. g**. Enrichment plot for the cardiac muscle contraction (KEGG) at 5 dpi (left panel) and heatmaps indicating expression of the leading-edge genes in mock or CHIKV-infected mice at 2 dpi or 5 dpi. N=5 mice/group. **h**. Enrichment plot for the viral arrhythmogenic right ventricular cardiomyopathy (KEGG) at 5 dpi (left panel) and heatmaps indicating expression of the leading-edge genes in mock or CHIKV-infected mice at 2 dpi or 5 dpi. N=5 mice/group. **i**. Enrichment plot for the viral myocarditis pathway (KEGG) at 5 dpi (left panel) and heatmaps indicating expression of the leading-edge genes in mock or CHIKV-infected mice at 2 dpi or 5 dpi. N=5 mice/group. **(e-h)**. NES = normalized enrichment score; grey shading indicates the genes belonging to the leading edge. P values were calculated using mixed-effects model test using Mann-Whitney test **(d-f)**.

**Extended Data Fig. 4.**
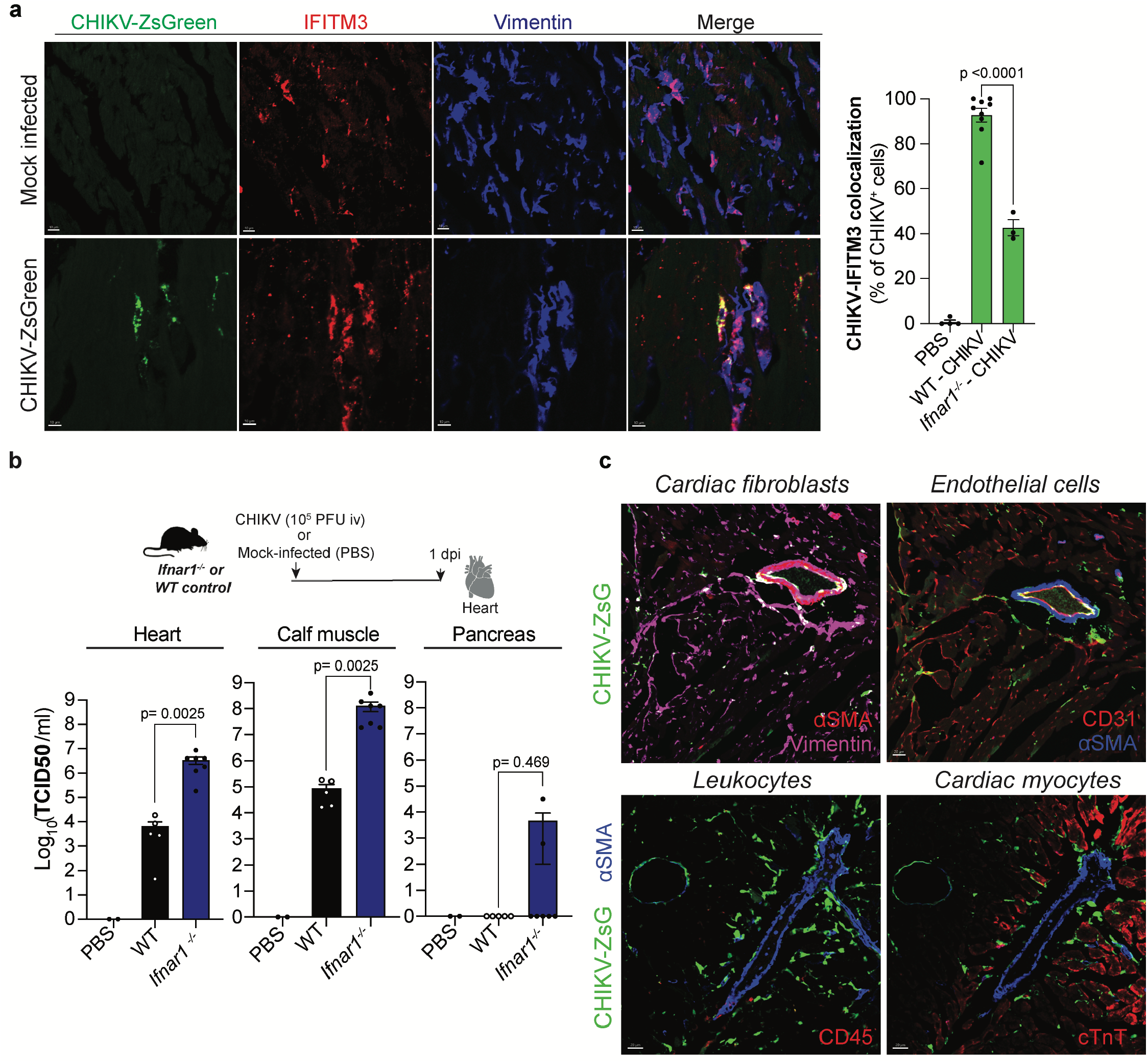
(Related to Fig.3). **a**. WT and *Ifnar1*^*-/-*^mice were inoculated intravenously with 1E5 PFU of reporter CHIKV-ZsGreen or mock-infected with PBS, harvested at 2 dpi, and cardiac tissue was stained with anti-IFITM3 and anti-vimentin antibodies. Left panel: Representative fluorescence microscopy images of ventricular sections of CHIKV-infected hearts and PBS control. Scale = 30 μm. Right panel: Co-localization analysis between CHIKV-infected cells and IFITM3 marker for WT and *Ifnar1*^*-/-*^ infected or mock-infected mice. Data are represented as the percentage of the total infected cells overlapping with the IFITM3 maker. Colocalization analysis was done on independent sections for N=2-3 mice/group **b**. Isolation of CHIKV infectious particles from heart, calf muscle and pancreas of WT or *Ifnar1*^-/-^ mice. Mice were infected intravenously with 1E5 PFU of CHIKV or PBS. CHIKV infectious particles were determined by TCID_50_ in BHK-21 cells. Mock-infected (N= 2), CHIKV-infected WT, or *Ifnar1*^-/-^ mice (N= 5-7 mice/group). **c**. Representative fluorescence microscopy images of ventricular sections of *Ifnar1*^-/-^ infected hearts (scale = 30 μm) colocalizing with markers of cardiac fibroblast, endothelial cells, leukocytes, and cardiac myocytes. **d**. P values were calculated using Mann-Whitney test **(a**,**b)**.

**Extended Data Fig. 5.**
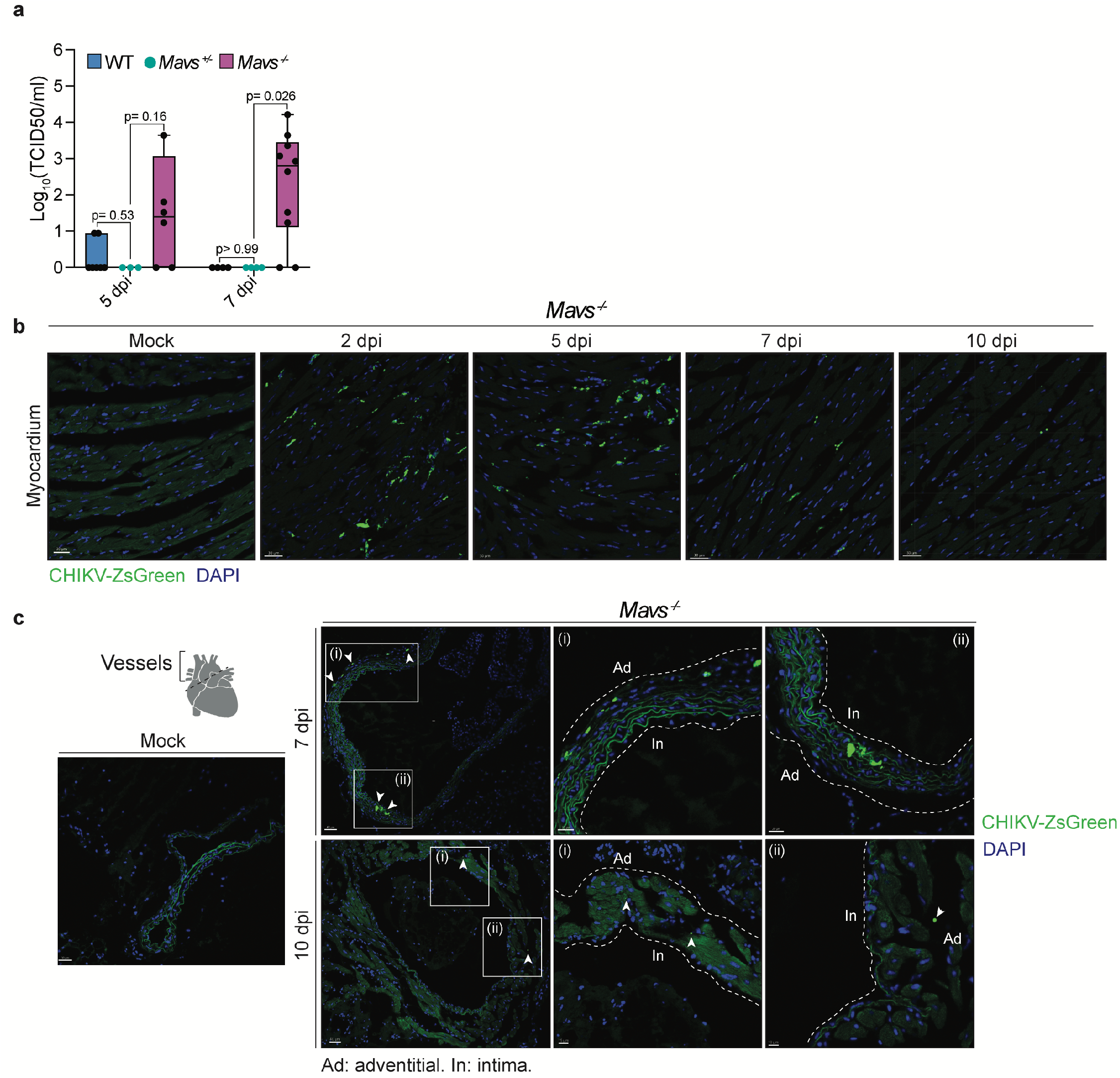
(Related to Fig.4). **a**. CHIKV cardiac infection kinetics in *Mavs*^-/-^, *Mavs*^+/-^ and WT mice. Mice were infected intravenously with 1E5 PFU of CHIKV. CHIKV infectious particles from heart homogenates were determined by TCID_50_ in BHK-21. N=3-10 mice/group. Data set from *Mavs*^-/-^ and WT mice are the same from **Fig 4b** at time point 5 and 7 dpi. **b**. Representative fluorescent microscopy images of ventricular sections of CHIKV-ZsGreen infected *Mavs*^-/-^ hearts. Scale=30 μm. **c**. Representative fluorescent microscopy images of vessels of CHIKV-ZsGreen infected *Mavs*^-/-^ hearts. Scale= 10-40 μm. P values were calculated using multiple Mann-Whitney tests.

**Extended Data Fig. 6.**
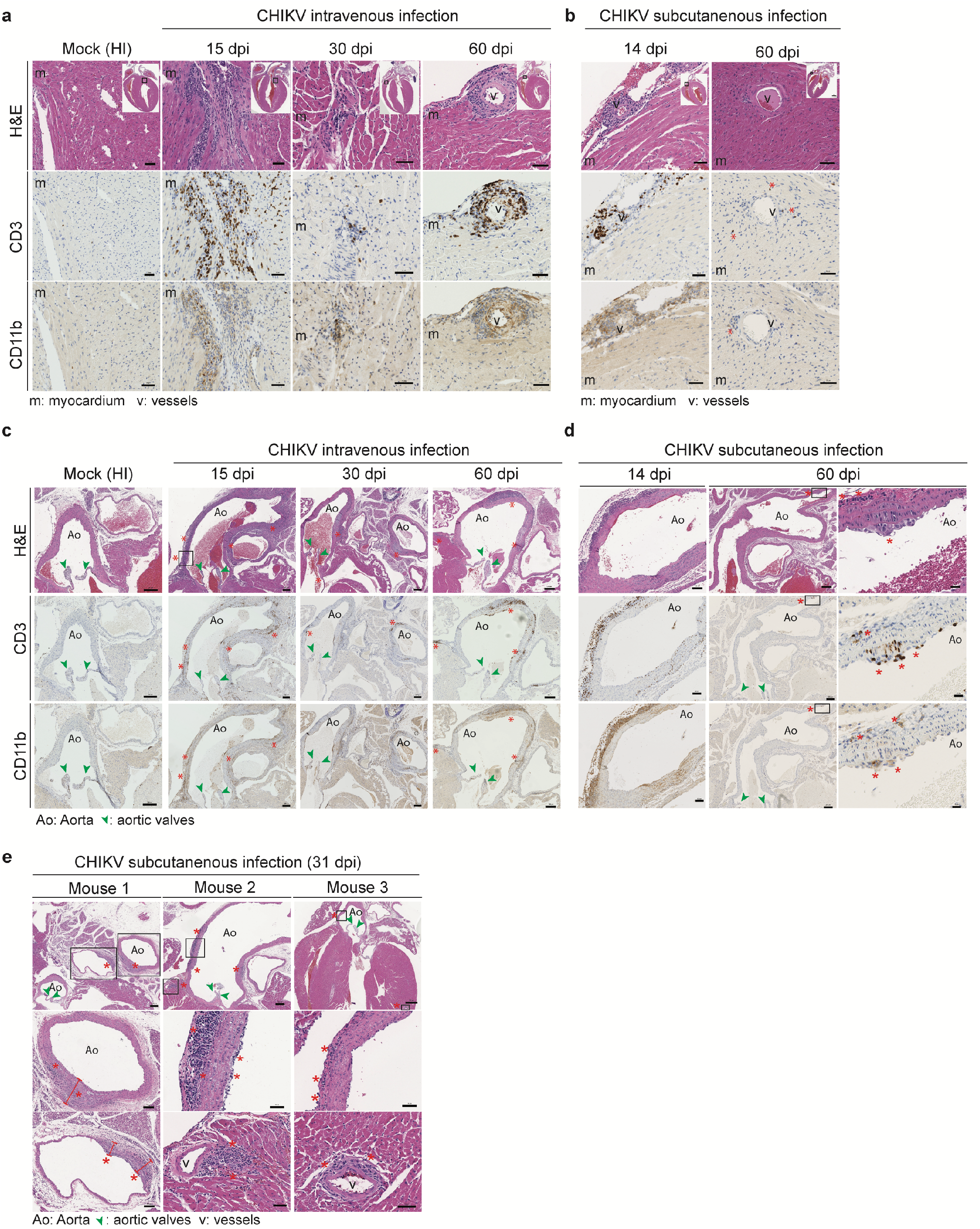
**a and c**. Representative H&E, CD3 and, CD11b staining for intravenously inoculated CHIKV or mock-infected *Mavs*^*-/-*^ at 15 dpi, 30 dpi, and 60 dpi, for the myocardial region **(a)** or for the major vessels attached to the base of the heart **(c)**. Inset shows the four-chamber view representation. **b and d**. Representative H&E, CD3, and, CD11b staining for subcutaneously inoculated CHIKV or mock-infected *Mavs*^*-/-*^ at 14 dpi, and 60 dpi, for the myocardial region **(b)** or for the major vessels attached to the base of the heart **(d)**. Inset shows the four-chamber view representation. **e**. Representatives H&E staining for subcutaneously inoculated CHIKV *Mavs*^*-/-*^ at 31 dpi. Scale bar is indicated at the bottom of each figure.

**Extended Data Fig. 7.**
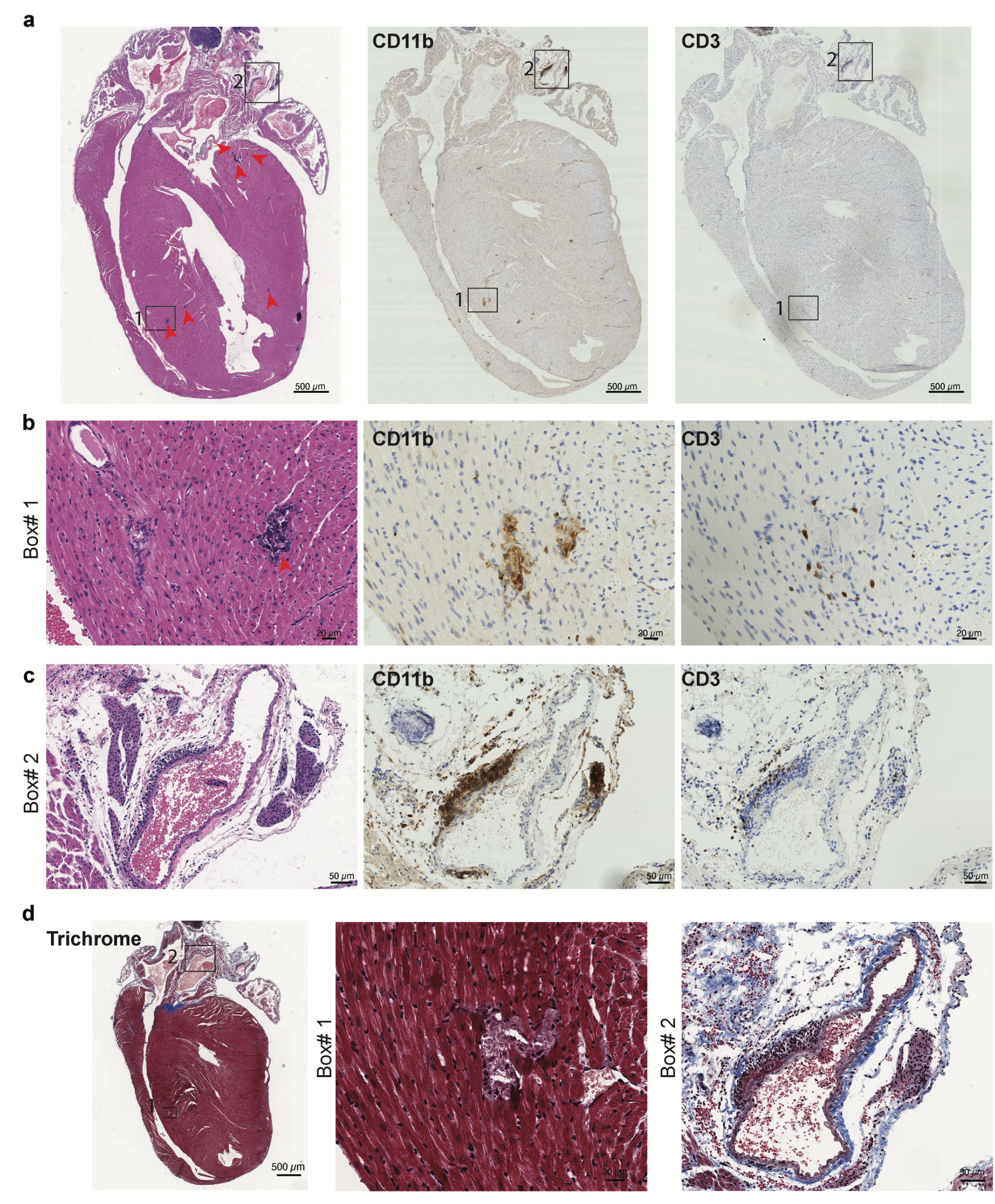
**a**. Four-chamber view representation of cardiac tissue from a 15 dpi *Mavs*^*-/-*^ mouse that succumb to CHIKV infection **(See Extended Data Table 1)**. Left panel, H&E showing four-chamber view and major vessels at the base of the heart. Middle panel: consecutive section stained with CD11b antibody. Right panel: consecutive section stained with CD3 antibody. Scale = 500 μm. **b**. Magnification of the ventricular region section (Box #1) for H&E, CD11b and CD3 staining showing infiltrates. Scale = 20 μm. **c**. Magnification of the pulmonary artery (PA) section (Box #2) for H&E, CD11b and CD3 staining. Scale = 50 μm. **d**. Mason Trichrome staining, showing no signs of fibrosis.

